# Protein compactness and interaction valency define the architecture of a biomolecular condensate across scales

**DOI:** 10.1101/2022.02.18.481017

**Authors:** Anton A. Polyansky, Laura D. Gallego, Roman G. Efremov, Alwin Köhler, Bojan Zagrovic

**Author notes:** equal contribution (Anton A. Polyansky, Laura D. Gallego). To whom correspondence should be directed:.

## Abstract

Non-membrane-bound biomolecular condensates have been proposed to represent an important mode of subcellular organization in diverse biological settings. However, the fundamental principles governing the spatial organization and dynamics of condensates at the atomistic level remain unclear. The *S. cerevisiae* Lge1 protein is required for histone H2B ubiquitination and its N-terminal intrinsically disordered fragment (Lge1_1-80_) undergoes robust phase separation. This study connects single- and multi-chain all-atom molecular dynamics simulations of Lge1_1-80_ with the *in vitro* behavior of Lge1_1-80_ condensates. Analysis of modelled protein-protein interactions elucidates the key determinants of Lge1_1-80_ condensate formation and links configurational entropy, valency and compactness of proteins inside the condensates. A newly derived analytical formalism, related to colloid fractal cluster formation, describes condensate architecture across length scales as a function of protein valency and compactness. In particular, the formalism provides an atomistically resolved model of Lge1_1-80_ condensates on the scale of hundreds of nanometers starting from individual protein conformers captured in simulations. The simulation-derived fractal dimensions of condensates of Lge1_1-80_ and its mutants agree with their *in vitro* morphologies. The presented framework enables a multiscale description of biomolecular condensates and embeds their study in a wider context of colloid self-organization.

## INTRODUCTION

Biomolecular condensates, such as P-bodies, nucleoli and stress granules, are membrane-less structures that contribute to the compartmentalization of the cell interior (Brangwynne et al., 2009; Feng, Chen, Wu, & Zhang, 2019; Lafontaine, Riback, Bascetin, & Brangwynne, 2021; Mitrea & Kriwacki, 2016; Mittag & Pappu, 2022; Molliex et al., 2015). They have been implicated in diverse biological functions including transcription, signaling and ribosome biogenesis (Banani, Lee, Hyman, & Rosen, 2017; Boeynaems et al., 2018) and have also been linked with different pathologies (Alberti & Dormann, 2019). Importantly, major efforts have been invested in understanding the general physicochemical principles behind the formation of biomolecular condensates (Alberti & Hyman, 2021; Banani et al., 2017; Brady et al., 2017; Brangwynne, Tompa, & Pappu, 2015; Bremer et al., 2022; Dignon, Zheng, Best, Kim, & Mittal, 2018; Martin et al., 2020; Mitrea & Kriwacki, 2016; Pappu, Cohen, Dar, Farag, & Kar, 2023; Zeng, Ruff, & Pappu, 2022). While an early paradigm for understanding condensate formation has been liquid-liquid phase separation, mounting evidence suggests that a close coupling between segregative phase separation and associative network transition, i.e., percolation, may be important in many cases (Choi, Holehouse, & Pappu, 2020; Choi, Hyman, & Pappu, 2020; Mittag & Pappu, 2022; Pappu et al., 2023; Schmit, Bouchard, Martin, & Mittag, 2020; Seim et al., 2022)

Formation of biomolecular condensates has been observed for proteins (Nott et al., 2015; Wang et al., 2018; Wei et al., 2017), DNA (J. T. King & Shakya, 2021), RNA (Jain & Vale, 2017) and their mixtures (Garcia-Jove et al., 2019). Intrinsically disordered proteins (IDPs), in particular, feature extensively in many condensates and are thought to contribute to their formation via transient intermolecular contacts (Banani et al., 2017; Uversky, 2021). An important challenge in this regard has been to provide a multiscale picture of the spatial organization and dynamics of IDP condensates, connecting the conformational properties and interaction patterns of individual polypeptides in a crowded environment with the features of the condensates they build. Considering that IDP condensates are extremely dynamic and structurally heterogeneous, it is clear that such a picture needs to capture the statistical, ensemble-level aspects of how matter inside the condensates is organized.

While an atomistic view of individual proteins in crowded environments is currently beyond the reach of high-resolution experimental techniques, molecular dynamics (MD) simulations have been developed with precisely this aim in mind (Dror, Dirks, Grossman, Xu, & Shaw, 2012). Although limited in terms of sampling efficiency as compared to coarse-grained approaches (Benayad, von Bülow, Stelzl, & Hummer, 2021; Dignon et al., 2018; Martin et al., 2020), atomistic MD simulations can provide an accurate picture of protein structure, dynamics and interactions with sub-Ångstrom resolution. For example, atomistic simulations have been used to model the dynamics of different peptides such as elastin-like peptide (Rauscher & Pomes, 2017) or different fragments of NDDX4 (Paloni, Bailly, Ciandrini, & Barducci, 2020) in condensate environment. Such simulations have also been combined with coarse-grained approaches and experiments (Zheng et al., 2020) and have provided a detailed view of the key interactions behind protein condensate formation (Conicella et al., 2020; Murthy et al., 2019; Ryan et al., 2018). Despite these important advances, however, the study of biomolecular condensates at the atomistic resolution is still at its beginning and many questions related to their most generalizable features are only starting to be addressed (Conicella et al., 2020; Li, Casalini, Arosio, & Salvalaglio, 2022; Murthy et al., 2019; Ryan et al., 2018). These, in particular, concern the complex interplay between structure, dynamics and thermodynamics of biomolecules in crowded environments.

In general, the concepts and methods of polymer physics have been widely used to understand the formation of biomolecular condensates (Alberti & Hyman, 2021; Brady et al., 2017; Brangwynne et al., 2015; Dignon et al., 2018; Dzuricky, Rogers, Shahid, Cremer, & Chilkoti, 2020; Guillén-Boixet et al., 2020; Martin et al., 2020; Mitrea & Kriwacki, 2016; Pappu et al., 2023). However, in contrast to the dense, continuous phase observed upon macroscopic phase separation in simple polymers, many protein condensates tend to have a lower density and be enriched in water (Alberti & Hyman, 2021; Keating & Pappu, 2021; Wei et al., 2017; Zaslavsky & Uversky, 2018), making them closer in organization to typical colloids (Slomkowski et al., 2011). According to the definition used in colloidal chemistry, biological condensates share features with weak gels, which undergo a transition between a population of finite-size pre-percolation clusters or sol, and an infinitely large cluster or gel (Stauffer, Coniglio, & Adam, 1982). In the case of biological condensates, phase separation coupled to percolation results in finite-sized colloidal clusters and the appearance of surface tension. In particular, under the requirement that the saturation concentration (c_sat_) for phase separation is lower than the percolation concentration (c_perc_), phase separation leads to an increase in local protein concentration and defines the phase boundary, while a percolation transition establishes network connectivity (Mittag & Pappu, 2022).

Importantly, starting with the seminal work of Forrest and Witten (Forrest & Witten, 1979), it has been recognized that aggregates or clusters of colloidal particles typically exhibit fractal scaling, i.e. that the cluster mass scales with cluster size according to a non-integer power law with the so-called fractal dimension *d_f_* as the exponent (Lazzari, Nicoud, Jaquet, Lattuada, & Morbidelli, 2016). In contrast to the regular geometric fractals, colloids generally build *statistical fractals* in which scaling laws hold between *average* values of mass and cluster size (Havlin & Ben-Avraham, 1987; Stanley, 1984). Overall, statistical fractal properties have been demonstrated under different conditions for many non-biological colloidal systems including silica, polystyrene and gold colloids (Lazzari et al., 2016). Moreover, by using scattering techniques such as static and dynamic light scattering (Lazzari et al., 2016; M. Y. Lin et al., 1989), or confocal and scanning electron microscopy (Khatun et al., 2020), the fractal nature has been associated with several biological colloidal systems, including different protein fibrils (Knowles et al., 2007; Nicoud, Lazzari, Balderas Barragán, & Morbidelli, 2015), whey protein isolates (Kharlamova, Nicolai, & Chassenieux, 2020) and clusters of lysozyme (da Silva & Arêas, 2005) and amylin (Khatun et al., 2020). Finally, fractal model has recently been used to interpret the results of coarse-grained simulations and characterize the aggregation of a phase-separating intrinsically disordered huntingtin fragment (Ruff, Khan, & Pappu, 2014).

Following the paradigm set by fractal colloidal systems (Lazzari et al., 2016), we derive here a general analytical framework for modelling statistical fractal cluster formation involving biomolecules. As a key result, we connect the interaction valency and compactness of individual biomolecules in condensates with the fractal dimension *d_f_*, which in turn enables us to describe the structural organization of the condensate at an arbitrarily chosen length scale. We apply the above framework to the N-terminal 80-residue fragment of Lge1, a scaffolding protein required for histone H2B ubiquitination during transcription (Gallego et al., 2020). Lge1_1-80_ exhibits a strong compositional bias shared by many known phase-separating proteins (enriched in Y, R and G, Figure 1A), is fully disordered and undergoes phase separation readily (Gallego et al., 2020), making it a powerful system to study the general features of condensate formation in IDPs. We combine a detailed atomistic characterization of a model condensate containing 24 copies of Lge1_1-80_, obtained via microsecond MD simulations (Figure S1A), with our newly developed fractal formalism in order to propose an atomistic model of the Lge1_1-80_ condensate extending to micrometer scale. To probe sequence determinants of Lge1_1-80_ LLPS, we investigate its all-R-to-K (R>K) mutant, where the ability to form condensates *in vitro* is modulated, and its all-Y-to-A (Y>A) mutant, which impairs phase separation *in vitro* (Gallego et al., 2020) (Figure 1B). Finally, we compare our theoretical predictions against the phase behavior of these different systems as studied experimentally via light microscopy. Our analyses provide a detailed, multiscale model of a biologically relevant condensate and establish a general, experimentally testable framework for studying other similar systems.

**Figure 1.**
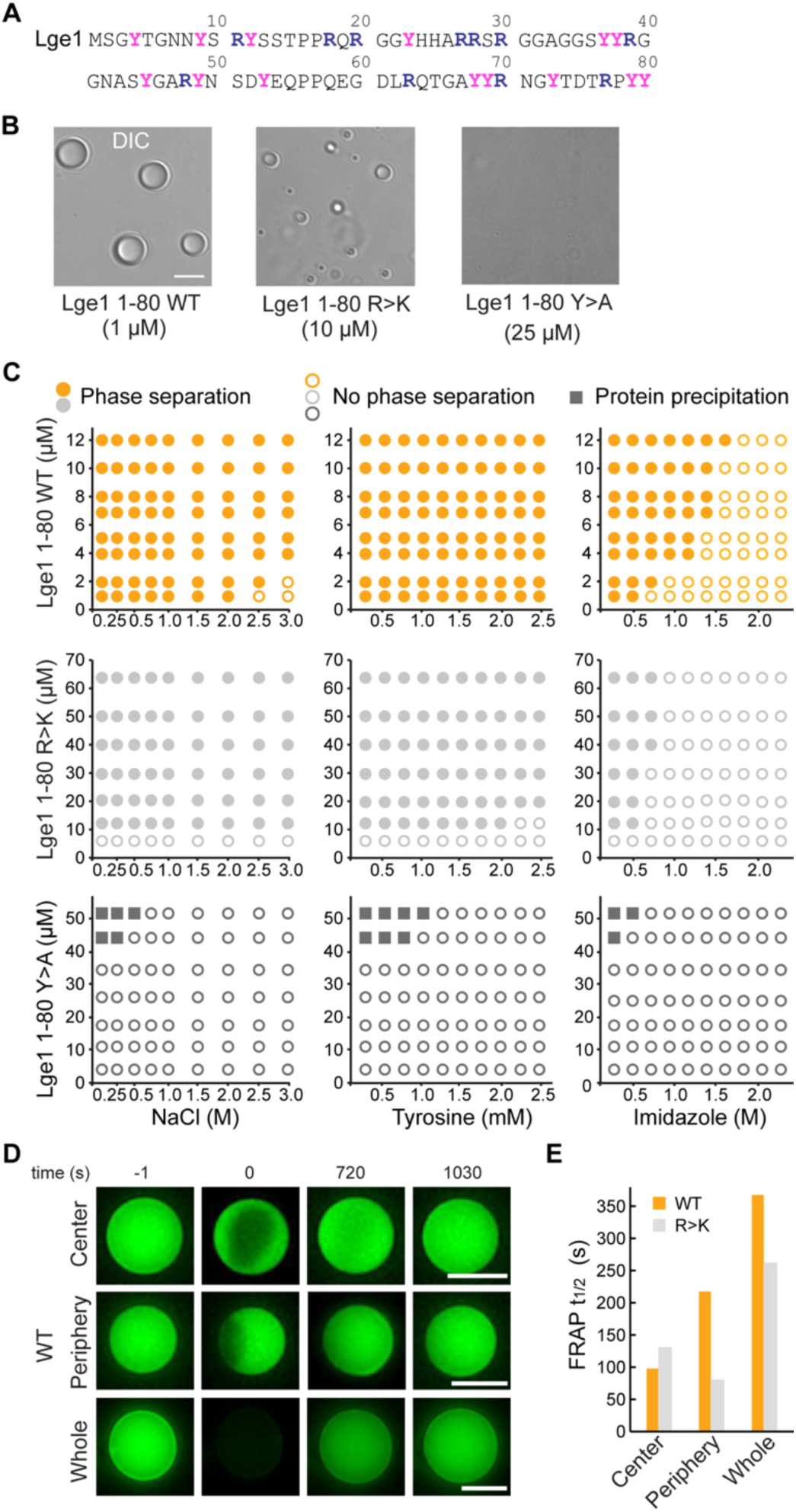
Lge1_1-80_ LLPS critically depends on tyrosine residues. (**A**) Sequence of Lge1_1-80_. Arginines and tyrosines are highlighted in deep blue and magenta, respectively. (**B**) Condensate formation for Lge1_1-80_ WT (left) and R>K (middle) in buffer with 200 mM NaCl. No such condensates are observed for Lge1_1-80_ Y> A (right). Scale bar, 5 µm. (**C**) Solubility diagrams for the WT (*top row*), R>K (*middle row*) and Y>A (*bottom row*) Lge1_1-80_ variants with protein concentration given on the *y*-axis and concentration of NaCl (*left panels*), tyrosine (*middle panels*) and imidazole (*right panels*) given on the *x*-axis. (**D**) Representative FRAP images of Lge1_1-80_ WT condensates, bleached in the center (*upper panels*), periphery (*middle*) or across the whole condensate (*lower panels*), including pre-bleach (*left, time -1s*), bleach (*time 0 s*) and post-bleach (*time 720 s, 1030 s*). Scale bars, 5 µm. (**E**) Half-times of fluorescent recovery after photobleaching (FRAP) of Dylight-labelled Lge1_1-80_ WT (orange) and R>K (grey) that were bleached in the center, periphery or across the whole condensate. Data was obtained after fitting to a double exponential model (see Figure S2).

## RESULTS

### Lge1_1-80_ LLPS critically depends on tyrosine residues

We have first experimentally explored the solubility diagrams of WT Lge1_1-80_ and its R>K and Y>A mutants by light microscopy using recombinant Lge1_1-80_ peptides (Figure S1B). WT Lge1_1-80_ phase separates already at a protein concentration of 1 µM in 0.1 M NaCl (Figure 1B, C). Its phase separation is robust and can be inhibited at high salt concentration (>2 M NaCl). The presence of tyrosine amino acids, which could provide competing π-π interactions, has no significant impact on WT condensate formation in the concentration range tested, while imidazole disrupts it at concentrations above 0.5 M. Mutating all arginine residues in WT Lge1_1-80_ to lysine (R>K mutant) increases the c_sat_ for phase separation by approximately 10-fold under the conditions tested (Figure 1B, C). The droplet-like WT and R>K condensates (Figure 1B, S1C) display normal fusion behavior and exhibit high circularity (Figure S1D). On the other hand, the substitution of the arginine guanidium groups, which are prone to π-π interactions, by the lysine amino groups, which lack sp2 electrons and do not participate in π-π interactions, makes the R>K mutant more sensitive to aromatic agents such as tyrosine (moderate effect) or imidazole (strong effect) (Figure 1C). At the same time, Y>A mutations strongly impair phase separation (Figure 1B, C, S1C, see also (Gallego et al., 2020), Ext. Data Figure 5), suggesting that tyrosines play a critical role in WT and R>K phase separation (Figure 1C) and highlighting the importance of π-π interactions in biomolecular condensate formation, as proposed earlier (Vernon et al., 2018). Moreover, the solubility diagrams are also in line with the observation that phase separation is related to the number of tyrosine and arginine residues in FUS and other phase-separating proteins (Bremer et al., 2022; Dzuricky et al., 2020; Wang et al., 2018). Finally, we note that at protein concentration of 45 μM and above, the Y>A mutant results in sporadic amorphous precipitates (Figure 1C).

We have used fluorescence recovery after photobleaching (FRAP) to characterize the overall dynamics of the condensates formed by WT and R>K Lge1_1-80_ variants. First, the applied FRAP protocol was tested for LAF1 (see Methods) and a fluorescence recovery of up to 80% in approximately 20 min was demonstrated (Figure S2H, I), in agreement with the previously published data (Taylor, Wei, Stone, & Brangwynne, 2019). While the Y>A mutant was shown to be unable to form droplet-like clusters, the dynamic WT and R>K condensates were studied using different bleaching strategies, including bleaching at the center, the periphery or across the whole condensate (Figure 1D, Figure S2A-G). Calculation of the recovery half-times by using different single exponential models did not provide quality fits (S2 source table, Figure S2H). Using double-exponent fitting allowed us to improve the quality of the fits and accurately describe the data (Figure 1E, Figure S2A-C, and S2 source table). Thus, the WT and R>K recovery half-times are in the range of 100 s for partial bleaching, and increase up to 350 s for bleaching of the whole condensates under the conditions tested. These values are similar to those previously reported for the *in vitro* condensates of other phase-separating proteins (Y. Lin, Protter, Rosen, & Parker, 2015; Taylor et al., 2019). Importantly, the recovery half-time for the whole bleached WT condensates is approximately 30% higher than for R>K (Figure 1E), suggesting a potentially different internal organization of the two types of condensates.

To rationalize the biochemical effects of the mutations, we have calculated the interaction strengths of selected pairwise contacts (Y-Y, R-Y, K-Y). To this end, the binding free energies (ΔG) for these pairs sidechain analogs were determined using all-atom Monte-Carlo simulations in chloroform, methanol, DMSO and water separately (Figure S1E, Figure S1E source table). Interestingly, ΔG(Y-Y) is independent of the polarity of the environment (approx. -5 kcal/mol on average), while ΔG(R-Y) and ΔG(K-Y) depend significantly on the dielectric permittivity of the medium (Figure S1E), as shown for other aromatic sidechain analogs (F, W) (Polyansky, Volynsky, Arseniev, & Efremov, 2009) and nucleobases (de Ruiter, Polyansky, & Zagrovic, 2017) interacting with R and K. Overall, both ΔG(R-Y) and ΔG(K-Y) are lowest in the apolar environment (approx. -10 kcal/mol on average), intermediate in bulk water (approx. -7.5 kcal/mol on average) and highest at intermediate polarity, where they are comparable to ΔG(Y-Y). The strong dependence of ΔG(R-Y) on the properties of the environment and the fact that, regardless of the conditions, ΔG(Y-Y) is never more favorable than ΔG(R-Y) were also recently reported for simulations of complete amino acids (Krainer et al., 2021), although the exact values are difficult to compare due to differences in the exact nature of the simulated systems, the force fields used and the method of how the ΔGs were derived.

In our simulations, significantly stronger R-Y binding as compared to K-Y was observed only in the environments of intermediate polarity (Figure S1E). This could contribute to the observed increase in the concentration required for Lge1_1-80_ R>K phase separation as compared to the WT, especially if Lge1 condensates exhibit a lower dielectric constant than that of bulk water, as observed elsewhere (Nott et al., 2015). The effect of R>K substitutions on phase-separation behavior was also examined by others for different IDPs (Bremer et al., 2022; Dzuricky et al., 2020; Schuster et al., 2020; Wang et al., 2018) and was, furthermore, linked with the general differences in the intrinsic physicochemical properties of R and K (Dubreuil, Matalon, & Levy, 2019; Fisher & Elbaum-Garfinkle, 2020; Fossat, Zeng, & Pappu, 2021; Hong et al., 2022; Paloni, Bussi, & Barducci, 2021; Zeng et al., 2022). Our aim here is to explore it further in the context of Lge1_1-80_ single- and multi-chain protein-protein interactions. Finally, the Y>A substitution likely causes a strong thermodynamic effect due to the removal of all possible R-Y and Y-Y intermolecular contacts.

### π-π interactions shape protein clustering in Lge1_1-80_ condensates

To better understand the effect of mutations on the organization of Lge1 condensates, we have performed three 1-µs-long MD simulations with 24 copies each of Lge1_1-80_ WT, Y>A, and R>K mutants in the mM concentration range as well as control simulations of the three proteins present as single copies (Figure S1A for details). In our simulations, proteins tend to form clusters that are characterized by pronounced structural heterogeneity and dynamics (Figure 2A). Specifically, the WT 24-copy system forms a single percolating cluster over the last 0.3 µs (Figure 2C) with all copies of the protein engaged. On the other hand, the mutants do not form such an extensive interaction network, but rather associate in multiple, differently sized clusters. The large continuous WT cluster is shaped by π-π interactions (Figure 2A, B) between the most abundant amino acids (R, Y, G), whereby R-Y contacts dominate (10 % of all possible pairs, Figure S3A), followed closely by G-Y and Y-Y contacts (7 % of all possible pairs each, Figure S3A). In particular, both R-Y and Y-Y contacts are strongly enriched over the sequence background (see Methods for definition), especially in the intermolecular context, i.e. between different protein chains in multi-chain WT simulations (Figure S3A, B). While glycine residues contribute significantly to the intermolecular interactions in WT with high absolute frequencies of G-Y and G-R contacts (Figure 2B), these contacts are either only slightly enriched over the expected sequence background (G-Y) or are even significantly depleted (G-R), as indicated in Figure 2B source table. The latter suggests that in WT R and Y prefer to contact residues in the sequence other than G, which is also the case for single-chain interactions (Figure S3A, B).

**Figure 2.**
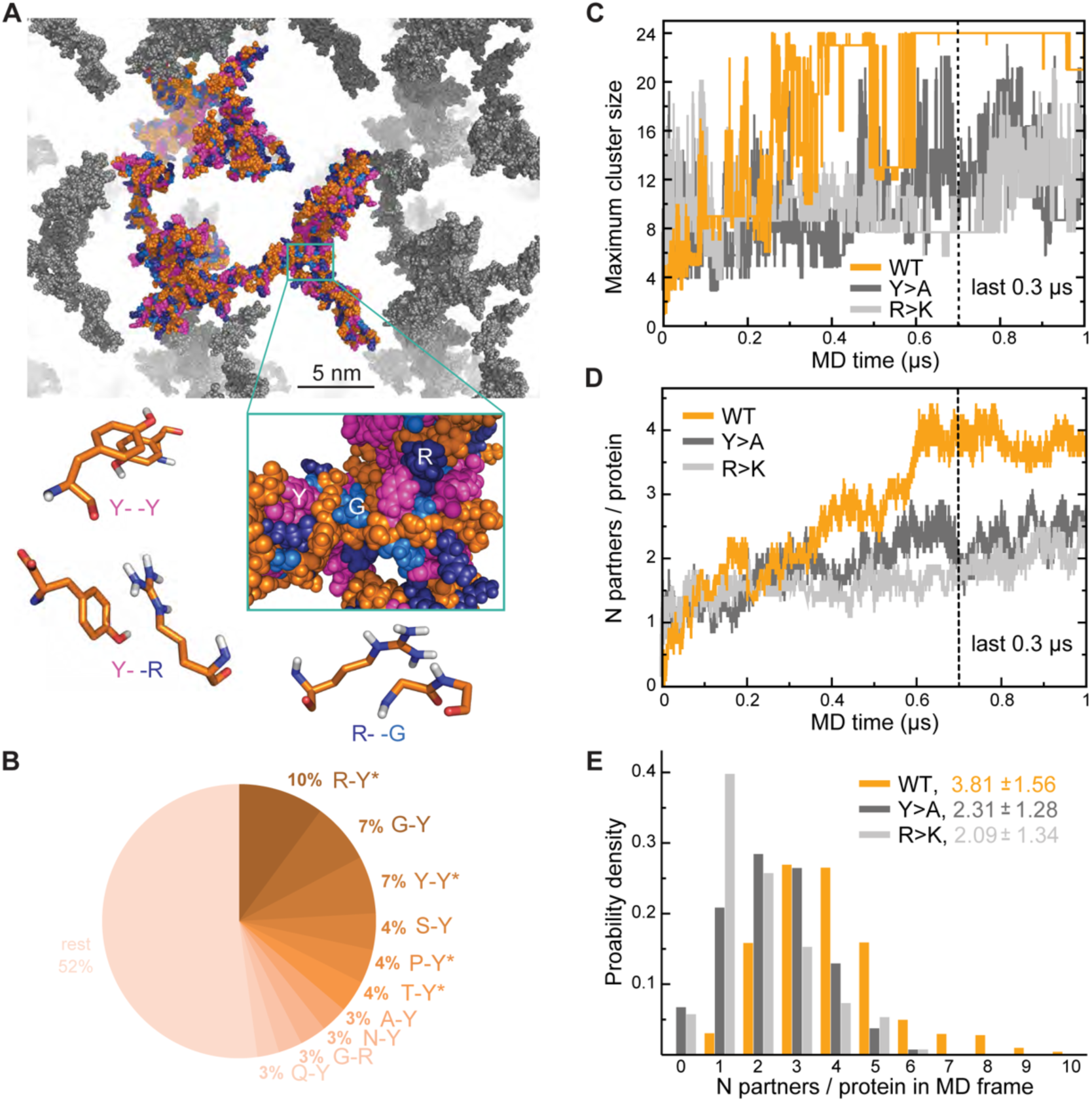
Analysis of interaction networks for Lge1_1-80_ variants by all-atom MD in 24-copy systems. (**A**) Exemplary MD-snapshot of the WT interaction network. Proteins in the simulation box given in the atomic representation (orange), whereby glycine, arginine and tyrosine residues are colored in sky blue, deep blue and magenta, respectively. Periodic images of the simulated system are shown in gray. (**B**) Pairwise contact statistics over the last 0.3 µs of simulations for the Lge1_1-80_ WT. Contacts enriched over the sequence background are marked with stars (see Fig S3A for exact enrichment values). (**C**) Time evolution of the largest detected protein cluster size and (**D**) the average number of interaction partners per protein chain, i.e. interaction valency, for the WT (*orange line*), Y>A (*dark gray line*), and R>K (*light gray line*) multi-chain systems. (**E**) Distributions of the number of interaction partners per protein over the last 0.3 µs of simulations. The color code is the same as in (**C**) and (**D**).

Interestingly, R>K substitutions increase the importance of G-Y contacts with respect to multi-chain interactions, where they rank top and are even more enriched than K-Y contacts. At the same time, for both WT and R>K Lge1_1-80_ variants the Y-Y contacts are enriched with respect to intramolecular interactions in single-chain simulations and become even more enriched in intermolecular interactions in multi-chain ones (Figure S3A). However, the opposite is seen for other homotypic contacts such as R-R in WT and Y>A or G-G in all three variants, since these contacts are enriched only for intramolecular interactions in single-chain simulations and are depleted for intermolecular interactions in multi-chain ones (Figure S3B). Altogether, our results show that R-Y is the top contact type in WT and together with Y-Y drives the multi-chain association of this IDR, in agreement with recent observations (Bremer et al., 2022). The results also point at a specific set of contacts which undergo significant rewiring when going from intramolecular interactions in single-chain simulations to intermolecular interactions in crowded systems, as further discussed below.

### Lge1_1-80_ interaction valency is defined by its sequence composition

Interaction valency, defined as the average number of binding partners per protein molecule, stabilizes in the course of WT simulations and reaches the average value of approximately 4 over the last 0.3 µs (Figure 2D), with individual WT copies having anywhere between 1 and 9 partners at some point (Figure 2E). In contrast, multiple non-interacting protein copies are still observed for the two mutants throughout the simulations, with the average valencies plateauing around 2 and the maximum number of partners not exceeding 6 in either case (Figure 2E). Thus, mutations with impaired phase separation as detected experimentally exhibit a significantly lower valency of protein-protein interactions in our simulations (see Technical summary table for details). Differences in valency can be also translated into different probabilities of contact formation, a key concept in percolation theory. We have estimated contact probabilities from simulations under the assumption of a well-mixed system, i.e. that all chains in the simulation box can in principle establish contacts with all the other chains. As shown in Figure S3F, contact probabilities evolve in direct proportion to valency, with a plateau over the last 0.3 µs. Notably, the WT contact probability reaches a level that is ∼1.5 fold higher than for either mutant. This contact probability is sufficient to form a single percolating cluster with all 24 copies interconnected, i.e. is higher than the critical contact probability at which one expects the network transition, which is not seen for the two Lge1_1-80_ mutants (Figure 2C).

The interactions between IDPs in our simulations are characterized by a *dynamic binding mode* where the interacting sequence motifs (“stickers”) and the non-interacting sequence motifs (“spacers”) (Martin et al., 2020) can be described only statistically, and no well-defined, specific structural organization of protein complexes is detected (Figure 3A). For WT, the typical binding mode, which corresponds to the average valency of 4, is characterized by different partners binding along the sequence in multiple, highly interactive regions (Figure 3A, B). In line with valency decrease, both mutants display fewer prominent interaction motifs, whereby for the phase separation-disruptive Y>A mutant intermolecular binding is detected for the C-terminal third of the molecule only (Figure 3A).

**Figure 3.**
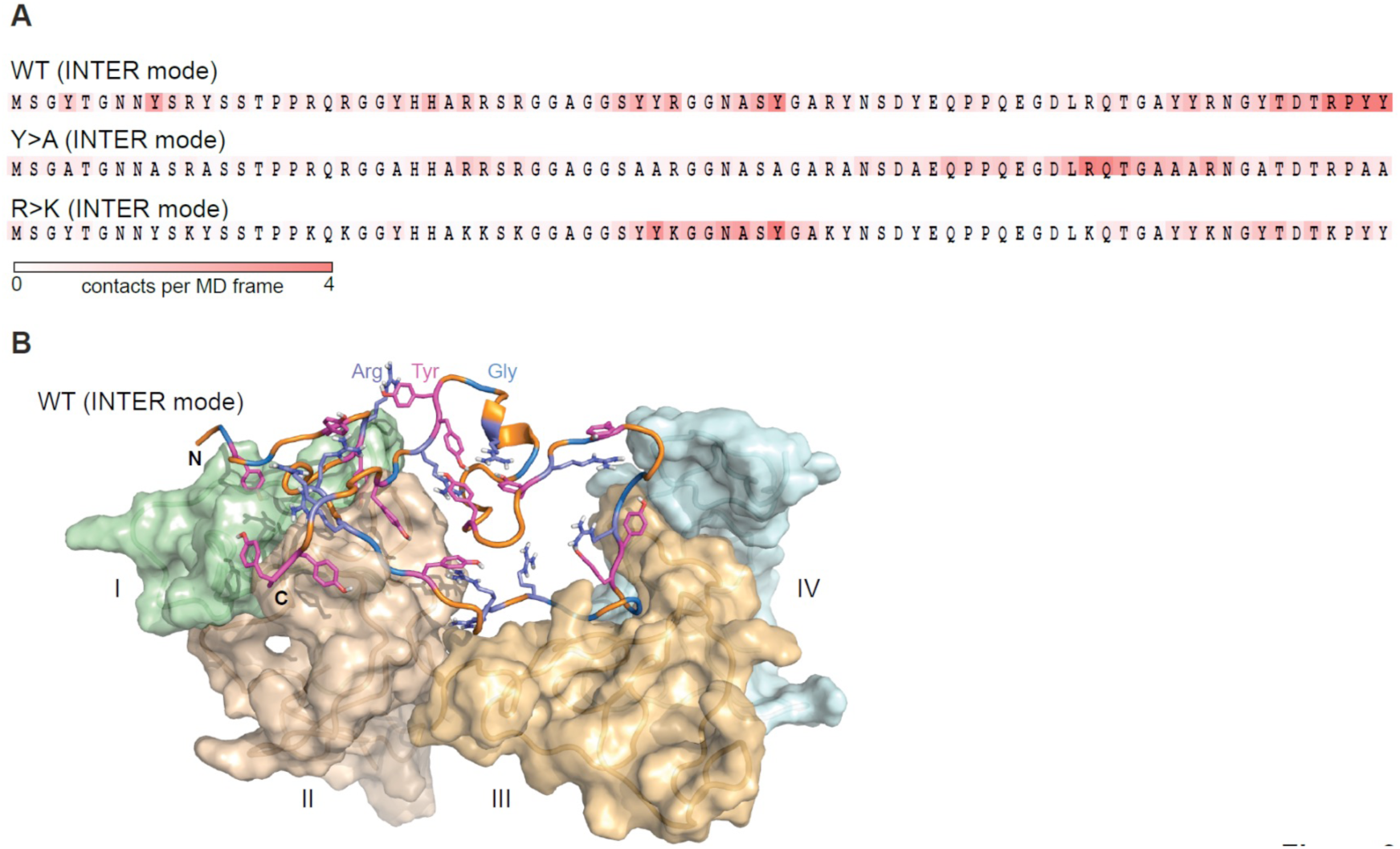
Lge1_1-80_ variants exhibit a dynamic binding mode in multi-chain systems. (**A**) Representative distributions of statistically defined interaction regions (“stickers”) mapped onto the protein sequence. Protein sequences are colored according to the average contact statistics over the last 0.3 µs. Four interaction profiles of proteins having the number of partners corresponding to the average valency in the system (4 partners for the Lge1_1-80_ WT and 2 partners for both mutants) and displaying the highest mutual correlations were used for the determination of the representative mode. The representative modes in all three cases display the Pearson correlation coefficient *R* > 0.6 with the interaction profiles obtained by averaging over all 24 copies in each system. (**B**) 3D model of the representative binding mode for Lge1_1-80_ WT. An MD snapshot at 1 µs is given for a protein copy (shown in cartoon and sticks representation; the color scheme is the same as in Fig. 2B) that is simultaneously interacting with four partners (shown in surface representation in pale cyan, pale green, wheat and light orange, and indicated by Roman numerals).

Interestingly, the dynamic modes identified in the intramolecular context are clearly different for all modeled proteins (Figure S3C), which is also partially observed at the level of pairwise contact statistics (Figure S2A, B), as discussed above. Specifically, intra- and intermolecular interactions rely on a similar pool of contacts by amino-acid type, but differ significantly if one analyzes specific sequence location of the interacting residues involved (Figure S3A and B). For example, one observes a high correlation between the frequencies of different contacts by amino-acid type when comparing intramolecular contacts in single-chain simulations and intermolecular contacts in multi-chain simulations (Figure S3D). This correlation is completely lost if one analyzes position-resolved statistics (2D pairwise contacts maps) or statistically defined interaction modes (Figure 3A, and S3C, S3E). Interestingly, the core of intramolecular interactions observed for a single molecule at infinite dilution and in the crowded context remain approximately the same, as reflected in the high correlation between intramolecular modes obtained in single- and multi-chain simulations (Figure S3E). The latter suggests that proteins maintain core self-contacts and establish new ones with neighbors, but do not lose self-identity as expected in a polymer melt. The observed ‘symmetry breaking’ between intra- and intermolecular mode of IDRs interactions is in line with the recent study by Bremer et al. (Bremer et al., 2022) and Martin et al. (Martin et al., 2020). In particular, Lge_1-80_ exhibits a relatively high net charge per residue (0.075), a non-uniform patterning of tyrosines (Ωaro=0.47, p=0.57, see Martin et al. for methodological details) and a high abundance of arginines, all of which could contribute to symmetry breaking, as proposed in these studies.

### Lge1_1-80_ sequence impacts its conformational behavior and dynamics

The perturbation of intra- and intermolecular interaction networks by mutations results in a different conformational behavior of the resulting Lge1_1-80_ variants. At the level of single molecules, extensive interactions involving R and Y residues result in a substantial compaction of the WT chain with <*Rg_MD_*> = 1.58 ± 0.12 nm (Figure 4A, see also Technical summary table for details). In contrast, the *Rg_MD_* distributions of the R>K and Y>A variants cover a wider range and display significantly higher average values (1.81 ± 0.35 and 1.71 ± 0.29 nm, respectively, Figure 4A). This is in all three cases more compact as compared with the predictions for a random coil of the same length (<*Rg_rc_>* = 2.50 nm, (Bernado & Blackledge, 2009)). On the other hand, the difference between WT and the two mutants is directly related to the extent of intramolecular interactions: the looser the interaction network, i.e. the fewer long-range sequence contacts there are, the larger the <*Rg>* (Figure S4A). In the crowded environment, WT again adopts a compact organization with an almost invariant <*Rg_MD_>* = 1.60 ± 0.22 nm when averaged over all 24 copies (Figure 4B), while the weakly self-interacting Y>A mutant unwinds toward more extended conformations (<*Rg_MD_>* = 1.97 ± 0.43 nm). At the same time, R>K, being most loosely packed in the single-molecule context, comes closer to WT values in the crowded environment (<*Rg_MD_>* = 1.67 ± 0.27 nm, Figure 4B). The latter can potentially be explained by the repulsive nature of K-K contacts, which are enriched in the single-molecule context and depleted in the crowded phase (Figure S3B). Note here that, unlike K-K pairs, R-R pairs can engage in π-π interactions as reflected in relatively higher fractions and enrichments of these contacts for Y>A, particularly in the single-molecule context (Figure S3B), as previously observed in PDB structures (Vernon et al., 2018). These results highlight the fact that the conformational behavior of IDPs in the bulk or in the crowded phase displays a clear sequence-specific character and cannot easily be generalized.

**Figure 4.**
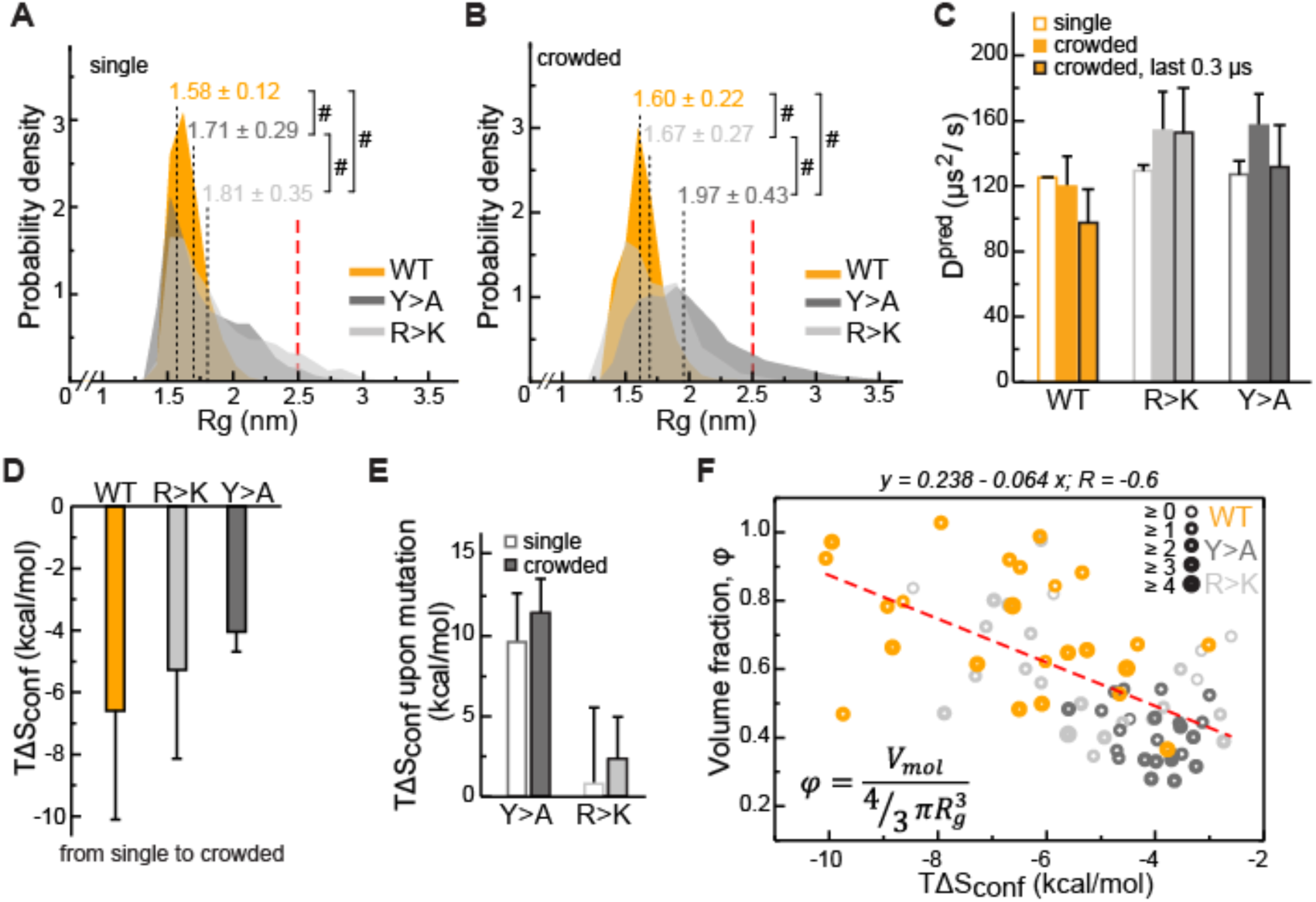
Impact of Lge1_1-80_ sequence on its conformational behavior and dynamics. Distributions of radii of gyration (*Rg*) for Lge1_1-80_ variants in (**A**) single-chain and (**B**) multi-chain systems. The last 0.3 µs of MD trajectories were used to collect *Rg* statistics for (**A**) 2 independent runs of the single-chain simulation and (**B**) all 24 protein copies in the multi-chain system. Average *Rg* values (<*Rg_MD_>*) and the corresponding standard deviations over the last 0.3 µs are indicated. Theoretical *Rg_rc_* value for an 80-aa disordered protein chain (see Methods) is shown with a vertical red dashed line. #, p-value < 2.2 10^-16^ according to Wilcoxon rank sum test with continuity correction. (**C**) MD-derived single-molecule translational diffusion coefficients of Lge1_1-80_ variants. For single copies the values were averaged between the two independent MD runs. For 24 copy systems the value was averaged between all proteins. Error bars depict standard deviations. (**D**) Average changes in the configurational entropy (Δ*S_conf_*) of a protein molecule for the transition from the single-molecule context (dilute state) to the crowded environment. Entropy values are given in energy units (*T*Δ*S_conf_*, *T* = 310 K) and were obtained using complete 1 µs MD trajectories. Averaging was done for entropy differences in all possible combinations between two independent runs of single molecules and 24 protein copies in the crowded system. (**E**) Average changes in *T*Δ*S_conf_* upon different mutations (see Methods) in the single-molecule context and in the crowded environment. Averaging was done for entropy differences over all possible combinations (2x2 and 24x24, respectively). (**F**) Correlation between relative single molecule configurational entropy changes (*T*Δ*S_conf_*) and the corresponding average compactness (φ) values of all protein copies in the crowded system. The average valency of different protein copies is proportional to the thickness of the circles as given in the legend. Entropy values, average compactness and average valency values were calculated over the complete 1 µs MD trajectories.

Single-molecule translational diffusion coefficients of Lge1_1-80_ variants obtained from fitting of MSD curves with an applied finite-size PBC correction and solvent viscosity rescaling (see Methods for details) are ∼120 µm^2^/s for all three single-chain simulations or anywhere between 100-150 µm^2^/s for multi-chain simulations and different Lge1_1-80_ variants (Figure 4C and Source Data 4C). In comparison, the diffusion constant of the similarly sized ubiquitin (Rg =1.32 nm, 76 aa, 8.6 kDa) at the protein concentration of 8.6 mg/ml is 149 µs^2^/s (Altieri, Hinton, & Byrd, 1995), while that of GFP (Rg =2.8 nm, 238 aa, 27 kDa) at the concentration of 0.5-3 mg/ml is ∼90 µs^2^/s (Baum, Erdel, Wachsmuth, & Rippe, 2014). This suggests that the diffusional dynamics captured by our simulations may be realistic. The obtained viscosity values (see Methods for details) in single-chain simulations of the three Lge1_1-80_ variants (effective concentration of 2.3 mg/ml) are all similar to each other and are close to the calculated solvent viscosity for TIP4P-D water/0.1 M NaCl of 0.83 mPa*s (Figure S4F and Source Data 4C). In the crowded multi-chain systems (effective concentration of 6-7 mg/ml), the viscosity systematically increases by about 20% and is again similar for all three Lge1_1-80_ variants (Figure S4F and Source Data 4C). These calculated values are in the range of reported values for other similar systems, e.g. serum albumin (Gonçalves et al., 2016).

In both single- and multi-chain simulations, the WT translational diffusion coefficients are somewhat lower than for either mutant (Figure 4C, and Source data). This effect does not appear to be related to protein size (<*Rg_MD_>,* Figure 4B) or viscosity (Figure S4F), but may reflect protein slow-down due to more extensive interactions with partners, at least in the crowded environment (Figure 2D). For instance, the WT diffusion coefficient drops by approximately 20% over the last 0.3 µs of the trajectory (Figure 4C), which correlates with the formation of a single percolating cluster in the system (Figure 2C). At the same time, the R>K diffusion constant does not change during the multi-chain simulation and is similar to the single-chain one (Figure 4C), likely due to electrostatic repulsion in the crowded environment. For Y>A there is no clear trend, whereby the diffusion constant over the last 0.3 µs of multi-chain simulations is similar to the single-chain one, which may be related to its larger size as compared to other variants (Figure 4B).

We have next quantified the effect of R and Y mutations on the Lge1_1-80_ conformational dynamics by estimating the configurational entropy (*S_conf_*) and its changes in different contexts via Maximum Information Spanning Tree (MIST) formalism. Due to the representation of molecules in internal bond-angle-torsion coordinates (see Methods), MIST is well suited for unstructured protein ensembles, as shown before (Fleck, Polyansky, & Zagrovic, 2018). *S_conf_* displays a reasonable convergence between the individual replicas of the single-chain simulation on the 1-µs time scale (<0.1 kJ/mol/K), especially in the case of the weakly self-interacting Y>A mutant (Figure S4B). Interestingly, in the crowded multi-chain environment, we observe a significant decrease in *S_conf_* for all three variants, with the biggest change seen for WT and the smallest for Y>A (Figure 4D). These results suggest that the crowding involves a conformational reorganization of the molecules toward decreasing the available free volume, i.e. increasing their compactness (volume fraction) φ, defined here as the ratio between the van-der-Waals and the hydrodynamic volume of a molecule. Finally, there is a substantial increase in *S_conf_* associated with the Y>A mutation of the WT in both single- and multi-chain contexts, in contrast with the R>K mutation, where a much weaker effect is observed (Figure 4E).

In the crowded environment, φ reaches plateau values over the last 0.3 µs (Figure S4C) with the average value fluctuating within a 2-4% interval for different averaging blocks (Technical summary table). Importantly, an increase in φ in the crowded environment correlates directly with an unfavorable Δ*S_conf_* (Figure 4F, Pearson *R* is -0.6), an effect which can potentially be compensated for by the favorable enthalpy upon forming the extensive multivalent interaction network as observed for the WT. Conversely, Δ*S_conf_* correlates poorly with the average number of bound partners (valency; Pearson *R* = -0.3) and there is no correlation between the compactness φ, and the valency of interactions (not shown). Being mutually largely independent, these two characteristics of protein molecules – compactness and valency – are the key parameters describing the organization of the corresponding crowded phase, which generally reflect the entropic and the enthalpic contributions to self-organization, respectively.

### Describing condensate architecture via a fractal scaling model

To extrapolate the atomistic-level properties of the crowded protein phase to larger length scales, we modeled the assembly of phase-separated condensates as an iterative, fractal process (Figure 5A, see also Appendix). Colloid fractal models typically start with an Ansatz capturing the power-law dependence between mass and size across different scales (Carpineti & Giglio, 1992; Lazzari et al., 2016). For conceptual clarity and to demonstrate where the power-law dependence comes from, our derivation starts from a simple physical picture of associating clusters, and yields the known scaling relationship bottom-up. Thus, we assume that initially individual protein molecules, characterized by a given volume fraction φ, interact to form clusters of a given average valency *n*. In the next iteration, these clusters arrange into higher-order clusters, whereby the average valency of each cluster and the volume fraction occupied by the clusters formed in the previous iteration remain constant at all levels of organization. For instance, if a single protein molecule binds four other proteins (*n* = 4), this results in the second-iteration cluster consisting of 5 protein molecules (Figure 5A). In the next iteration, this 5-mer arranges with four other 5-mers into a new cluster with the same valency of 4, resulting in a larger cluster with 25 molecules. According to this model, the smaller clusters from iteration *i* − 1 always occupy the same volume fraction φ of the apparent volume of the larger cluster from iteration *i* (taken as 2/3 in Figure 5A). This scenario results in a simple fractal formalism whose benefit is that it yields exact solutions for different features of the clusters at each iteration (see Appendix). Thus, for each iteration *i*, the formalism returns the number of molecules (eq. 1), the apparent volume (*V*_i_, eq. 4), the size (*R*_i_, eq. 5), the mass (*M*_i_, eq. 8) and the effective molar concentration of molecules in the cluster (*C*_i_, eq. 6). While the above derivation for simplicity uses the constant volume fraction and valency at each iteration, we want to emphasize that in the case of structurally heterogeneous statistical fractal clusters, these parameters would be equivalent to their time- and ensemble-averaged counterparts.

**Figure 5.**
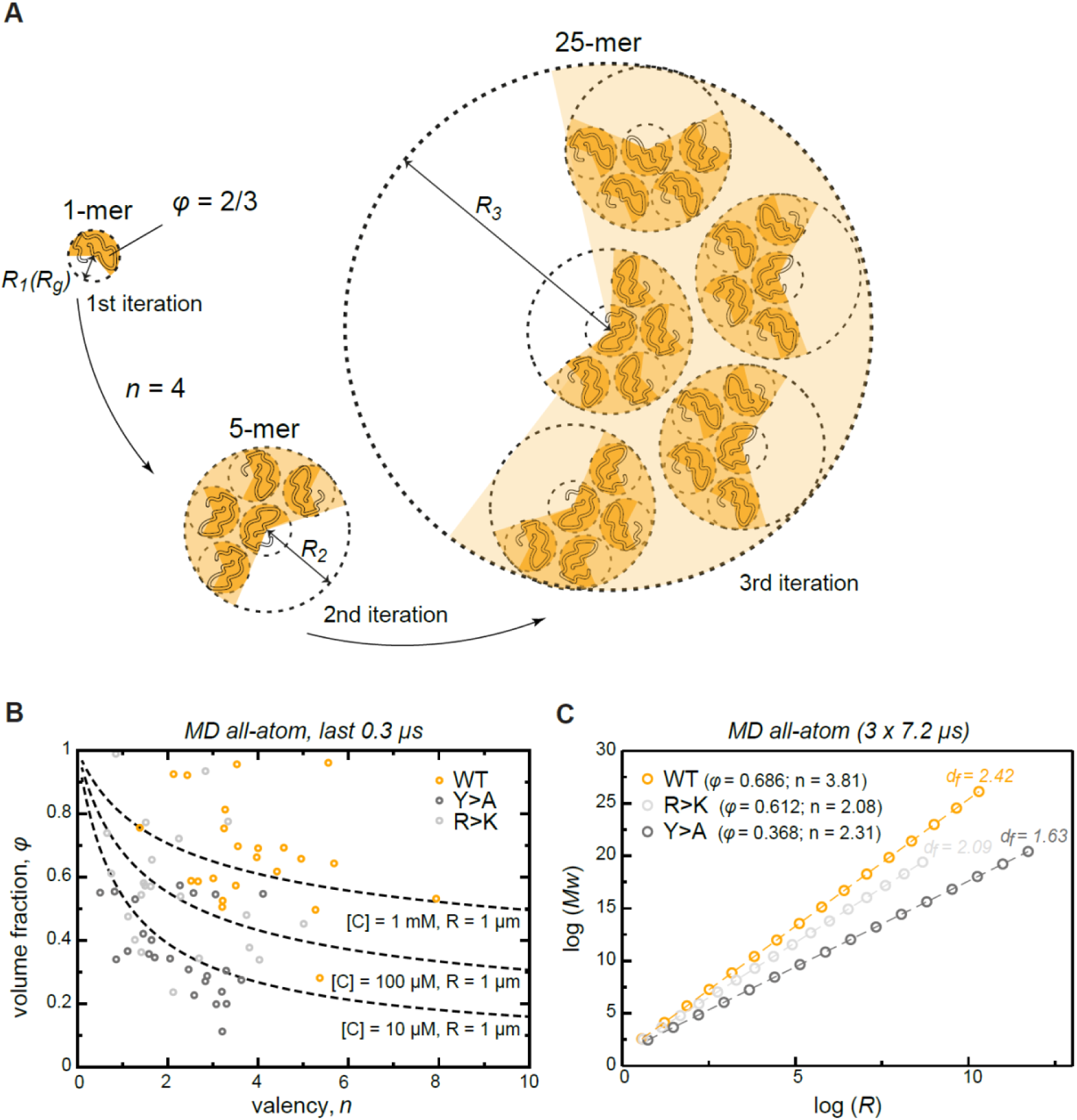
Describing condensate architecture via a fractal scaling model. (**A**) Schematic representation of a scaling principle in condensate assembly. Interaction valency (*n*) and compactness (*φ*) of individual proteins determine the properties of protein clusters at different iterations (see Appendix for the corresponding formalism). (**B**) The parameter space for *φ* and *n* for a condensate with a fixed size (R = 1 µm) and concentration ([C]) described by the model (*dashed lines*). The *φ* and *n* for each individual protein molecule in the crowded environment averaged over the last 0.3 µs of MD trajectories are shown with open circles. (**C**) Power-law dependence between mass and size of protein clusters at different iterations of the model with the applied valency and compactness corresponding to their average values over the 24 simulated protein copies and the last 0.3 µs of MD trajectories (indicated in the legend). Dashed lines show linear regression for the log *R* vs. log *Mw* plot with the corresponding slope or fractal dimension (*d_f_*) indicated above the lines.

We have compared the predictions of the fractal model with what is seen in the actual Lge1_1-80_ simulations. Importantly, the simulations in the first instance just give the average valency and compactness of individual chains in the dense phase. The fractal formalism, which is conceptually independent from the simulations, subsequently provides the dependence of condensate mass on its radius, M(R), at any desired length scale. This, in turn, enables one to directly test the predictions of the fractal formalism in the case of the actual clusters seen in the simulations. Thus, over the last 0.3 µs of MD simulations, the WT multi-chain system displays a narrow distribution of sizes for the single molecule (Figure 4B) and leads to a single percolating cluster (Figure 2D, S3C) with an average radius *R* = 6.27 ± 0.24 nm. The latter point agrees closely with the predictions of the fractal formalism when the average values for *φ* and *n* obtained over the last 0.3 µs of MD simulations are used (Figure S6). Thus, a single cluster consisting of 24 chains observed in simulation directly corresponds to the 3^rd^ iteration of the model (Figure S5A). Moreover, the slope (*A,* eq. 10) and the intercept (*B,* eq. 11) of the linear regression for the log *R* vs log *Mw* plots are also similar between the simulations and the model (2.33 vs 2.42 and 1.13 vs. 1.11, respectively) (Figure S5B). Finally, the latter parameters allow the exact calculation of the characteristic *φ* and *n* values (eq. 12 and 13; 0.638 and 3.76, respectively), which again are very similar to those obtained directly from all-atom MD (Figure S6). These non-trivial correspondences suggest that fractal organization is present even at the shortest scale, i.e. at the level of MD simulation boxes.

The model also enables one to explore the space of *φ* and *n* parameters for a condensate with a fixed size and protein concentration (eq. 7, Figure 5B). For instance, for a 1-µm condensate with a 1-mM apparent protein concentration, the corresponding *φ* and *n* values are generally in the range observed in MD for the crowded WT system. Generally, in order to keep the ratio of size-to-concentration fixed, protein compactness must decrease non-linearly with increasing valency (Figure 5B, dashed lines). Accordingly, perturbation of compactness and valency due to the two types of studied mutations can result in a decrease of the apparent concentration in condensates of fixed size. Importantly, the compactness of IDPs is not a stable parameter and is tunable by different factors (temperature, pH, ionic strength, etc) (Uversky, 2009). This, in turn, also suggests that IDP concentration inside condensates may also be adaptable and tunable. Thus, unwinding or compaction of an IDP due to any factor would change the apparent concentration and density in the condensates. The latter also opens up the possibility for potential microphase transitions inside of phase-separated droplets, in analogy to those known for lipid membranes (Lewis & McElhaney, 2013). Finally, protein concentration as a function of condensate size can be directly estimated from the model (Figure S5C). Notably, the complex topology of condensates (see also below) as proposed by the model allows for the formation of droplets with a very low apparent protein concentration.

### Valency and compactness define fractal dimension and condensate topology

The fractal model provides a direct relationship between size and mass of different clusters which directly capture the scaling behavior of the condensate matter. Thus, the slope of the line in the log R vs. log Mw plot (*A*) is equal to the fractal dimension *d_f_*, which describes the topology of molecular clusters (Carpineti & Giglio, 1992). The fractal dimensions equaling exactly 1, 2 or 3 correspond to the objects exhibiting 1D, 2D or 3D organization, respectively, while systems with non-integer *d_f_* have an intermediate dimensionality. The proposed model facilitates a direct investigation of the scaling properties in condensates for a molecule with defined characteristic compactness and valency of interactions. Most importantly, the fractal dimension of a condensate is completely defined by *φ* and *n* (eq. 10), reflecting the predictive potential of the proposed model. For instance, the average values of *φ* and *n* derived from MD simulations in the crowded state result in a different scaling behavior for WT, R>K and Y>A proteins, whereby their dimensionality respectively decreases (Figure 5C) and results in a different morphology of the corresponding condensates (Figure 6B), as observed experimentally (Figure 1B, C). Specifically, while the WT condensate simulated here exhibits a *d_f_* of 2.42 and is also experimentally found to undergo robust phase separation, the R>K mutant exhibits a lower *d_f_* of 2.09 and undergoes phase separation under a more limited set of conditions. Even more extremely, *d_f_* is 1.63 for the Y>A mutant (Figure 5C), which does not undergo phase separation and can be thought of as an object between 1D and 2D.

**Figure 6.**
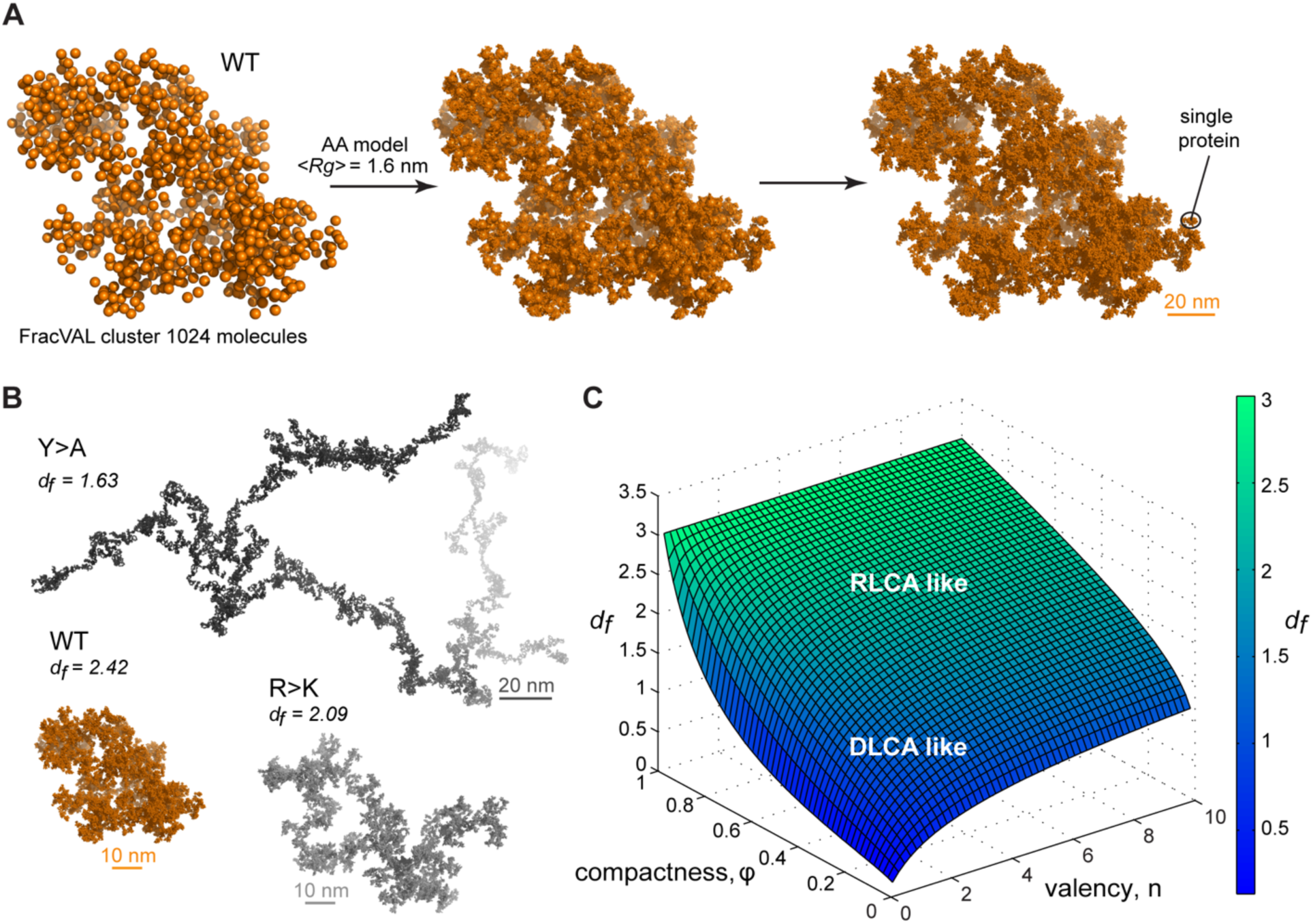
Reconstruction of the large-scale condensate architecture with atomistic resolution. (**A**) Transformation of a coarse-grained 1024 particle cluster obtained by FracVAL algorithm to an all-atom representation. The cluster was reconstructed using the fractal dimension *d_f_* and the averaged *Rg* value derived from multi-chain simulations of Lge1_1-80_ WT. (B) Representative 1024-protein clusters for Lge1_1-80_ variants at all-atom resolution. (**C**) The non-linear dependence of the fractal dimension *d_f_* on *φ* and *n* as given by the model formalism (see eq.10, Appendix). The surface is colored according to the corresponding *d_f_* values (see the scale bar).

### Reconstructing the condensate architecture across scales

There exist several algorithms in the colloid literature that enable one to reconstruct the geometry of fractal clusters starting from a given fractal dimension *d_f_* (Kätzel et al., 2008; Morán, Fuentes, Liu, & Yon, 2019; Thouy & Jullien, 1994) (and prefactor *k_f_*, which is equal 1 in the present model; see Appendix for derivation). Recently, Moran et al. (Morán et al., 2019) have proposed a robust and tunable algorithm, FracVAL, for modeling the formation of clusters consisting of polydisperse primary particles. We have used the values of *d_f_* derived from our simulations for the three Lge1_1-80_ variants in combination with FracVAL to generate individual realizations of the respective condensate structure on the length scale of hundreds of nanometers. In Figure 6A, we demonstrate this procedure for WT Lge1_1-80_: FracVAL produces cluster geometries using spherical particles with radii corresponding to the respective *<Rg_MD_>* values, which are then computationally replaced by the realistic protein conformations obtained from our simulations. In agreement with its micrometer-scale behavior observed *in vitro*, the modelled WT Lge1_1-80_ condensate exhibits a densely packed fractal geometry. In contrast, the reconstructed Y>A cluster exhibits an elongated, filamentous topology of low dimensionality (Figure 6B), which may preclude the formation of well-defined phase-separated droplets (Figure 1B and S1C). The R>K cluster exhibits an intermediate topology. The clusters shown in Figure 6B were generated using 1024 primary particles in all three cases: clearly, the Y>A cluster occupies a significantly larger volume as compared to the R>K cluster and especially the WT. In particular, the three reconstructed systems exhibit holes and cavities of different sizes, with Y>A being most extreme in this regard. Finally, note that the three reconstructions shown in Figure 6A and B are individual snapshots of the local architecture; the full fractal model entails an ensemble of such snapshots, all configurationally different, yet still conforming to the same scaling pattern.

## DISCUSSION

We have provided a multiscale description of Lge1_1-80_ condensates extending from a detailed analysis of constituent molecules at the atomistic level all the way to the micrometer-sized droplets involving thousands of individual molecules, which can be observed *in vitro*. We have shown that mutations of R and Y residues induce perturbations at the level of both intra- and intermolecular interaction networks, and result in conformational effects that can be related to the phase behavior and the ability to form condensates as detected by light microscopy. Specifically, the characteristic descriptors of protein behavior in the crowded phase, valency and compactness (volume fraction), are shown to be sufficient to describe the structural organization of condensates across length scales in the context of the proposed analytical fractal model. Importantly, the studied mutations substantially change either just the valency (R>K) or both the valency and compactness (Y>A) of Lge1_1-80_ polypeptides as shown in MD simulations.

The applied simulation protocol reproduces the level of diffusive protein dynamics expected from molecules of Lge1_1-80_ size. As a further indication of its general quality, the values of valency and compactness obtained from simulations are consistent with the difference in FRAP recovery dynamics observed for WT and R>K. Namely, the accurate fitting of FRAP data is possible only if using at least two components (Figure S2). According to (Sprague & McNally, 2005), these components reflect the contribution of particle diffusion and interactions. Thus, the recovery in centrally bleached condensates is faster for WT than for the R>K mutant, which can be related to the higher compactness of WT particle across scales, as compared to R>K (Figure 1E, S2A). On the other hand, the FRAP results for the condensates bleached in the peripheral area highlight the contribution of valency to condensate formation. Indeed, the recovery is about 3 times faster for the R>K mutant (Figure S2B), which could potentially be related to the lower valency of interactions and the ease of replacement of inactivated fluorescent species or/and exchange with proteins in the bulk. Indeed, a similar behavior with faster recovery for the R>K is observed when bleaching the whole condensate (Figure 1E, S2C).

In the fractal model, the changes in protein valency and compactness translate to different scaling behavior and, subsequently, different topology of the condensates. Brangwynne and coworkers have successfully adopted the theoretical formalism of patchy colloids to capture the relative contributions of oligomerization, RNA binding and structural disorder in the formation of stress granules and P-bodies (Sanders et al., 2020). In their analysis, they emphasized the role of the valency of colloid particles as a key parameter defining specificity and tunability of condensate features. Furthermore, Collepardo-Guevara and coworkers have used a coarse-grained patchy-particle colloid model to study the determinants of condensate stability and have highlighted the role of valency, while also clearly demonstrating the general impact of volume fraction (Espinosa et al., 2020). Our study now provides a general framework for assessing the importance of these two parameters on the formation of biomolecular condensates. Importantly, the proposed model predicts the existence and provides a quantitative characterization of topological properties of pre-percolation finite-size clusters that are in line with the recent findings (Kar et al., 2022; Mittag & Pappu, 2022; Pappu et al., 2023). More generally, the model provides the fractal dimension (*d_f_*) of protein clusters and enables evaluation of different scale-dependent properties of clusters of arbitrary size, including protein density as a function of cluster size (Figure S5C, Figure 5C). Finally, MD simulations of proteins in the crowded context on the length scale of tens of nanometers can be used in combination with cluster–cluster aggregation algorithms to derive atomistically resolved models of the 3D organization of fractal clusters of any chosen size (Figure 6A, B).

As discussed above, an emerging paradigm for biomolecular condensate formation is that of phase separation coupled to percolation. Importantly, fractal behavior, as explored in the present work, is naturally related to percolation phenomena. For instance, direct links between the fractal dimension defining scaling principles in self-similar clusters and critical exponents in different percolation models have been provided (Kapitulnik, Gefen, & Aharony, 1984; Stauffer et al., 1982). While a similar formal derivation for our model is out of the scope of the present study, we can provide an illustration of how our results and the key parameters of the model can be interpreted according to percolation theory. A key concept in percolation theory is that of contact probability between components in the system. The network transition appears and a percolating cluster is formed if the contact probability exceeds a particular threshold, i.e. the critical contact probability (*p_crit_*). For instance, in the simplest case according to the Flory-Stockmayer theory *p_crit_* = 1/(n-1), where n is a number of bonds formed by each monomer, and is related to the valency in our model. Thus, interaction valency contributes directly into the spatial organization of pre-percolating clusters (the fractal dimension) and defines the threshold of the percolation. Therefore, our analysis shows that WT Lge1_1-80_ displays a robust network transition due to high contact probability, which depends on its valency and, more generally, on the topology of its clusters. This perspective suggests that concentrations at which network transition is expected should be lower for WT than for either mutant, as indeed observed. Of note, the ability of IDRs to form low-dimensional fractal structures (*d_f_* is in a range of 1.6-1.9) upon the disruption of their tendency to phase separate by a polyalanine insertion was demonstrated experimentally for synthetic elastin-like polypeptides (Roberts et al., 2018).

The fractal formalism also provides a framework for approaching the question of protein concentration inside of phase-separated condensates, which covers a large range extending to tens of mM and beyond (Brady et al., 2017; McCall et al., 2020; Ryan et al., 2018). However, some proteins form condensates with extremely low concentrations in the dense phase. For example, the measured binodals of LAF-1 indicated that the concentration of the protein inside the droplet is 86.5 μM, which corresponds to an average separation between molecular centers of mass of 27 nm (Wei et al., 2017). Considering that the average *Rg* of LAF-1 is approximately 4 nm, it is not immediately clear how the molecules inside the droplet are organized in order to simultaneously establish intermolecular contacts and also form low-density droplets. Fractal organization provides a simple resolution of this apparent conundrum. Namely, fractal systems are characterized by a remarkable property that their packing density is a function of the length-scale on which it is examined (Stanley, 1984). For example, the density of a Sierpinski gasket, which is self-similar on all length scales, *decreases* exponentially with the length scale, with the exponent of *d_f_* - *d*, where *d* is the dimensionality of the space. Translated to the question of biomolecular condensates, fractal organization enables high local concentration of biomolecules at short length scales and, simultaneously, low global concentration at long length scales, as illustrated in Figure 5B and Figure S5C.

Importantly, fractal systems are characterized by the existence of holes, i.e. unoccupied regions of space, whose size covers a wide range of length scales (Stanley, 1984). The observation that the µm-sized WT Lge1_1-80_ droplets are fully permeable to dextrans *in vitro* (Gallego et al., 2020), even up to Mw = 2000 kDa or *Rg* ∼ 27 nm (Armstrong, Wenby, Meiselman, & Fisher, 2004) (Figure S3E, F), suggests that their organization allows for large holes (∼55 nm in diameter or more). This is consistent with our fractal model, which suggests that the dimensionality of WT condensates is below 3 (*d_f_* = 2.42, Figure 6). Finally, the multiscale nature of biomolecular condensates, as embodied in the statistical fractal model, also points to the possible formation of clusters with sizes well below the resolution limit of light microscopy. This may relate to some of the open questions in the condensate field, especially when it comes to their *in vivo* function (McSwiggen, Mir, Darzacq, & Tjian, 2019; Musacchio, 2022). Having said this, it should be emphasized that the proposed fractal model may be applicable to varying degrees in different systems. In particular, it was shown that *in vitro* Lge1 assembles a liquid-like core that is surrounded by an enzymatic outer shell formed by the E3 ubiquitin ligase Bre1 (Gallego et al., 2020). Whereas such condensates can exhibit a diameter of 1 µm or more when grown *in vitro*, they are expected to be smaller in cells (i.e. low nm range) (Gallego et al., 2020). Hence, the relevance of the proposed model will need to be tested experimentally with respect to nm-sized core-shell condensates in cells. Moreover, future studies must include the spherical Bre1 shell as a boundary condition which presumably constrains the 3D-orientation of the Lge1 C-terminus and thereby impinges on the geometry of the Lge1 meshwork.

Finally, the fractal model predicts the coexistence of differently sized clusters within a condensate, as reported recently (Kar et al., 2022), which have a characteristic scaling of mass with condensate size in the nm to µm range. This prediction of the model can be tested using static light scattering (SLS) techniques and will be a subject of our future work: a linearly decreasing intensity as a function of the scattering vector in a log-log representation, as frequently seen for different colloidal systems, is expected by the fractal model (Lazzari et al., 2016; M. Y. Lin et al., 1989). In fact, fractal dimension can be estimated from SLS experiments as the limiting value of scattering curves for high values of the product of the scattering vector *q* and the average cluster size *<RG>* (Hagiwara, Kumagai, & Nakamura, 1996). In addition, techniques such as DLS and MALS can be used in order to measure independently masses and sizes of LLPS condensates *in vitro.* It will be also important to analyze to what extent these features are retained in more complex, biologically relevant contexts.

Colloidal cluster formation is typically discussed in the context of two limiting regimes (Klein et al., 1990; Lazzari et al., 2016; M. Y. Lin et al., 1989). In diffusion-limited cluster aggregation (DLCA), the rate of cluster formation is determined by the time it takes for colloidal particles to encounter each other and every encounter leads to binding. In reaction-limited cluster aggregation (RLCA), particles need to overcome a repulsive barrier before binding and not every encounter is productive. Both regimes result in fractal behavior, with DLCA leading to looser structural organization and lower fractal dimensions and RLCA leading to more compact structural organization and higher fractal dimensions. Computer simulations and scattering experiments show that *d_f_* is ∼1.8 for DLCA and ∼2.1 for RLCA (Lazzari et al., 2016; M. Y. Lin et al., 1989). Importantly, Meakin and colleagues have shown that the two regimes of colloid cluster formation are universal and do not depend on the chemical nature of the underlying particles, making them an attractive paradigm for modelling the multiscale structure of biomolecular condensates as applied here (M. Y. Lin et al., 1989). However, for flexible, multivalent molecules like IDPs, the exact regime of cluster formation would depend on the values of *φ* and *n* and may be a tunable feature of the exact conditions. In Figure 6C, we present the non-linear relationship between the fractal dimension *d_f_* on *φ* and *n*, which exhibits certain general trends. For example, high values of either compactness *φ* or valency *n* or both, such as in the case of WT Lge1_1-80_, result in high values of *d_f_* and may be more associated with the RLCA model, while low values of both parameters may more be associated with the DLCA model (Figure 6C). A more quantitative analysis of the connection between the exact mechanism of cluster formation and the underlying parameters of compactness and valency will be a topic of future work.

Overall, our results provide an atomistic framework for understanding the role of valency and compactness of IDPs on condensate stability and architecture across scales. This presents an opportunity for the rational, quantitively founded design of phase-separating agents with predefined condensate properties. Indeed, recent studies have demonstrated the possibility to tune condensate properties *in vitro* and *in vivo* by manipulating the aromatic residue content and molecular weight in IDPs (Dzuricky et al., 2020) or the size of disordered linkers and valency in modular proteins (Lasker et al., 2021). Finally, it is our hope that our results may help to critically embed the field of biomolecular condensates in the wider context of colloid chemistry. We expect that the powerful theoretical, computational and experimental tools of colloid chemistry could propel the study of biomolecular condensates to the next level of fundamental understanding.

## METHODS

### Protein expression and purification

All proteins were expressed in *Escherichia coli* BL21 CodonPlus (DE3) RIL cells. 6His-Lge1 _1-80_-StrepII constructs were induced by addition of 0.5 mM isopropyl 1-thio-β-D-galactopyranoside at OD_600_ = 0.8 at 23 °C for 3 h and purified as published elsewhere (Gallego et al., 2020) with Talon® Superflow^TM^ beads (Cytiva) in a final elution buffer (10 mM Tris, 1 M NaCl, 1 mM TCEP, 1 M imidazole, 10 % vol/vol glycerol, pH 7.5) and stored at -80 °C. For protein labeling with Dylight^TM^ 488 NHS-Ester (Thermo Scientific), the final elution of the Lge1_1-80_ constructs was performed in 10 mM Hepes, 1 M NaCl, 1 mM TCEP, 1 M imidazole, 10% vol/vol glycerol, pH 7.5. Labelling was performed during the elution step for 45 min. Unbound dye was removed by sequential buffer exchange in centrifugal filters Amicon Ultra 0.5 ml 3 K (Merk Millipore). Lge1_1-80_-Dylight labelled protein was stored at -80 °C.

6His-LAF-1 was expressed and purified as described (Elbaum-Garfinkle et al., 2015) with some modifications as follows. Lysis buffer included 20 mM Hepes, 500 mM NaCl, 10% vol/vol glycerol, 14 mM β-mercaptoethanol, 10 mM imidazole, 1% vol/vol Triton 100, pH 7.5, and was supplemented with 0.5 mg/ml lysozyme, DNase I and protein inhibitor mix HP (Serva). After washing and eluting from Ni-NTA Sepharose 6 FastFlow beads (GE Healthcare) in elution buffer (20 mM Hepes, 1 M NaCl, 10% vol/vol glycerol, 14 mM β-mercaptoethanol, 250 mM imidazole, pH 7.5), protein was labelled with Dylight^TM^ 488 NHS-Ester (Thermo Scientific) for 30 min. Unbound dye was removed by sequential buffer exchange in centrifugal filters Amicon Ultra 0.5 ml 30 K (Merk Millipore). Finally, LAF-1-Dylight labelled protein was stored at -80 °C.

Protein quality was assessed by SDS-Page (4-12% gel, MOPS buffer) and Coomassie staining. Total purity of the protein was calculated by densitometry analysis of the gel bands with ImageJ. Percentage of the fraction of full-length protein was calculated in relation to all the bands present after purification.

### Solubility diagrams

Different concentrations of purified 6His-Lge1_1-80_-StrepII proteins in a total volume of 100 µL were added to a bottom-clear 96-well plate (Greiner Bio-One) in a buffer containing 25 mM Tris, 100 mM NaCl, 100 mM imidazole, 1% vol/vol glycerol and 1 mM DTT, pH 7.5. Varying concentrations of NaCl (100 mM to 3 M), imidazole (100 mM to 2.25 M) or tyrosine (0,25 mM to 2.5 mM) were analyzed as indicated in Figure 1C. Plates with the protein mix were incubated for 10 min at 20 °C. Turbidity of the samples was measured at 450 nm in a Victor Nivo™ plate reader (Perkin Elmer) at 20 °C. Assessment of protein phase separation or aggregation was performed by applying a total volume of 20 µL of the sample to a pretreated (Gallego et al., 2020) 16-well glass-bottom ChamberSLIP slide (Grace, BioLabs). DIC imaging was performed as previously described (Gallego et al., 2020).

### Circularity

Morphology of Dylight-labelled Lge1_1-80_ particles was assessed by studying circularity, calculated using the formula:

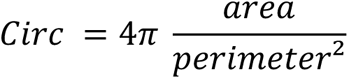

Image analysis was done in Fiji/ImageJ by applying the particle analyzer - shape descriptor plugin, and statistical analysis was conducted in GraphPad Prism v 7.0e. For each construct, at least four independent images at each respective protein concentration were analyzed (1 µM for WT and 10 µM for R>K). Total number of the particles analyzed (n) is included in the figure legend (Figure S1D).

### Fluorescent Recovery After Photobleaching (FRAP)

FRAP experiments were performed on a temperature controlled DeltaVision Elite microscope as previously described (Gallego et al., 2020). Lge1_1-80_ WT-Dylight condensates were formed by 100% labeled protein at a final concentration of 10 µM in 25 mM Tris pH 7.5, 100 mM NaCl, 1% vol/vol glycerol, 1mM TCEP, 100 mM imidazole. Lge1_1-80_ R>K-Dylight was mixed with 50% of unlabeled protein (given the enrichment in lysine residues that are labeled) to a final protein concentration of 30 µM. LAF-1-Dylight was mixed with 30% of unlabeled protein and processed as described (Elbaum-Garfinkle et al., 2015) to a final concentration of 8 µM in 25 mM Tris, pH 7.5, 100 mM NaCl, 1mM DTT. Bleaching was performed in protein condensates incubated for 30 min on pretreated 16-well glass-bottom slides (Gallego et al., 2020) by applying 20% power of a 50 mW laser 488 for 5 ms. Fluorescent intensity before bleaching was recorded for one frame prior to the bleach. Recovery of the bleach spot (central bleach, peripheral bleach or whole condensate bleach) was recorded elapsed in time to avoid photobleaching (initially every 7 s, then 14 s, 30 s and finally 60 s) with total 32 images for 20 min. Intensity traces were corrected for photobleaching. Recovery was calculated as published elsewhere (Taylor et al., 2019), normalized, and fitted to a double exponential function of the form:

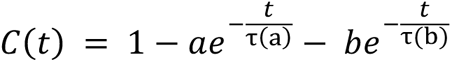

Finally, the recovery half times were obtained by the numerical solution of the fitting equation:

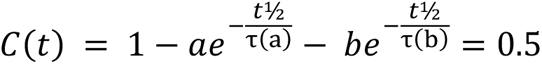

using WolframAlpha online (https://www.wolframalpha.com).

Total area of the bleached spot was calculated in ImageJ by relating it to the whole area of the condensate. Fluorescent intensity profiles for whole bleached condensates were acquired in ImageJ. For LAF-1, in addition to the double exponential function, several specific single exponent fits (Taylor et al., 2019) were tested in terms of quality of the fit.

### Dextran experiments

Experiments were performed as published elsewhere (Gallego et al., 2020). TRITC-Dextrans with a final concentration of 0.05 mg/ml were added to the samples containing 6His-Lge1_1-80_- StrepII (2 µM) in a final buffer containing 25 mM Tris, 100 mM NaCl, 100 mM imidazole, 1% vol/vol glycerol and 1 mM DTT, pH 7.5. Samples were incubated for 15 min at 20 °C on pretreated 16-well glass-bottom ChamberSLIP slides prior to imaging.

### Pairwise interaction free energy calculations

Pairwise interaction free energies were calculated using all-atom Monte Carlo (MC) simulations according to the previously established framework (Polyansky et al., 2009). MC simulations were carried out in TIP4P water (Jorgensen, Chandrasekhar, Madura, Impey, & Klein, 1983), methanol, dimethylsulfoxide and chloroform for the sidechain analogs of tyrosine, arginine, and lysine using OPLS force field (Jorgensen, Maxwell, & Tirado-Rives, 1996). Initial structures of the molecules were optimized *in vacuo* using AM1 semiempirical molecular orbital method (Dewar, Zoebisch, Healy, & Stewart, 1985) and placed in rectangular boxes with explicit solvent. All calculations were performed with the BOSS 4.2 program (Jorgensen & Tirado-Rives, 2005) with periodic boundary conditions in the NPT ensemble at 298 K and 1 bar. Standard procedures were employed including Metropolis criterion and preferential sampling for the solutes (Jorgensen & Ravimohan, 1985). The potential of mean force (pmf) computations for Y-Y, Y-K and Y-R pairs were performed by gradually moving the solute molecules apart in steps of 0.05 Å along an axis defined by the particular atoms or centers of geometry, while both solute molecules were allowed to rotate around this axis. Non-bonded interactions were truncated with spherical cutoffs of 12 Å. The free energy changes for a particular pair were calculated in a series of consecutive MC simulations using statistical perturbation theory and double-wide sampling (Jorgensen & Ravimohan, 1985). Each simulation consisted of 3 × 10^6^ configurations used for equilibration, followed by 6 × 10^6^ configurations used for averaging.

### Molecular dynamics (MD) simulations

The full-length structure of Lge1 was modeled *de novo* using Phyre2 web portal (Kelley, Mezulis, Yates, Wass, & Sternberg, 2015). The disordered N-terminal 1-80 aa fragment of Lge1 (Lge1_1-80_) in the modeled structure lacked any secondary structure and was used as an initial configuration in further Lge1 simulations. All-atom MD simulations were performed for WT Lge1_1-80_ fragment as well as for its Y>A and R>K mutants. Single-protein-copy MD simulations were carried out in 9 x 9 x 9 nm^3^ water boxes using 2 independent 1-µs replicas for each of the three Lge1_1-80_ variants. The same initial configuration (see above) was used for all three. All systems had zero net charge and effective NaCl concentration of 0.1 M. Systems with 24 protein copies were simulated in 18 x 18 x 18 nm^3^ (WT, Y>A) or 19 x 19 x 19 nm^3^ (R>K) water boxes. Initial configurations for these simulations were generated as follows. Four different protein conformers were selected from the initial 100 ns parts of the two independent single-chain MD simulations (two conformations from each run) based on the criteria of having been the centers of the most highly populated clusters after clustering analysis performed using *cluster* utility (GROMACS) with the applied RMSD cut-off for backbone atoms of neighboring structures of 1.5 Å. The cells containing 4 copies were assembled manually and translated 6 times in different directions, resulting in a protein grid containing 24 protein copies. A total of 1 µs of MD statistics were collected for each large system. All MD simulations and the analysis were performed using GROMACS 5.1.4 package (Abraham et al., 2015) and Amber99SB-ILDN force field (Lindorff-Larsen et al., 2010). After initial energy minimization, all systems were solvated in an explicit aqueous solvent using TIP4P-D water model (Piana, Donchev, Robustelli, & Shaw, 2015), which was optimized for simulating IDPs. The final NaCl concentration was 0.1 M (Figure S1A). The solvated systems were again energy-minimized and subjected to an MD equilibration of 30000 steps using a 0.5-fs time step with position restraints applied to all protein atoms (restraining force constants *Fx* = *Fy* = *Fz* = 1000 kJ mol^-1^ nm^-1^) and 250000 steps using a 1-fs time step without any restraints. Finally, production runs were carried out for all systems using a 2-fs time step. A twin-range (10/12 Å) spherical cut-off function was used to truncate van der Waals interactions. Electrostatic interactions were treated using the particle-mesh Ewald summation with a real space cutoff 12 and 1.2 Å grid with fourth-order spline interpolation. MD simulations were carried out using 3D periodic boundary conditions in the isothermal−isobaric (NPT) ensemble with an isotropic pressure of 1.013 bar and a constant temperature of 310 K. The pressure and temperature were controlled using Nose-Hoover thermostat (Hoover, 1985) and a Parrinello-Rahman barostat (Parrinello & Rahman, 1981) with 0.5 and 10 ps relaxation parameters, respectively, and a compressibility of 4.5 × 10^−5^ bar^−1^ for the barostat. Protein and solvent molecules were coupled to both thermostat and barostat separately. Bond lengths were constrained using LINCS (Hess, Bekker, Berendsen, & Fraaije, 1997).

Radii of gyrations (*Rg*) of simulated proteins were calculated using GROMACS *gyrate* utility, respectively. The average number of interaction partners per protein and the detailed statistics of intermolecular contacts were evaluated using GROMACS *mindist* and *pairdist* utilities with an applied distance cutoff of 3.5 Å, respectively, while intramolecular contact maps were generated using *mdmat* utility. Statistical significance of the difference between calculated parameters was evaluated using the Wilcoxon rank sum test with a continuity correction using *R* package (version 3.2.3). Protein structures were visualized using PyMol (Schrodinger & DeLano W, 2020).

### Pairwise MD contact statistics and the dynamic interaction mode

Frequencies of pairwise contacts between different positions in protein sequences were collected over the last 0.3 µs of MD trajectories independently for every simulated protein chain. These frequencies can be represented as position-resolved 2D maps or can be collapsed as total interaction preferences at each position along the sequence (1D interaction profile or *dynamic interaction mode*). Finally, they can be grouped by contact type and converted to pairwise frequencies and enrichments. An enrichment for a pairwise contact *A*-*B* is calculated as:

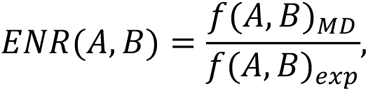

where *f*_*MD*_ is an observed MD frequency of contacts between *A* and *B* and *f*_*exp*_ is the frequency of such contacts expected at random, given the sequence composition of the chain i.e. *f*(*A*, *B*)_*exp*_ = *f*(*A*) × *f*(*B*), where *f*(*A*) is the frequency of X in the sequence. Individual 1D interaction profiles were obtained for each simulated protein considering only intramolecular protein contacts (INTRA) or only contacts with partners (INTER). Four individual interaction profiles of proteins having the number of partners corresponding to the average valency in the system over the last 0.3 µs and displaying the highest mutual correlations were used for the determination of the representative INTER mode. Interaction profiles averaged between the individual MD replicas of single-chain simulations were used for the determination of a representative INTRA mode.

### Theoretical estimate of Rg

The estimation was done according to the scaling model used in the polymer theory to connect *Rg* and the chain length of a random coil as follows:

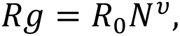

where N is the length of Lge1_1-80_ (*N* = 80) and *R*_/_ and *ν* are the empirical parameters refined for IDPs (*R*_/_ = 0.254 nm; *ν* = 0.522) (Bernado & Blackledge, 2009).

### Cluster analysis

The largest protein-protein interaction clusters in the 24-copy simulated system were identified using hierarchical clustering. For this purpose, minimum-distance matrices were calculated from each MD trajectory sampled at every 100 ps using GROMACS *mindist.* The clustering was done in MATLAB (R2009) using function *cluster* with an applied distance cutoff of 3.5 Å.

### Entropy calculations

The configurational entropy was evaluated by applying the maximum information spanning tree (MIST) approximation (B. M. King, Silver, & Tidor, 2012) using the PARENT suite (Fleck, Polyansky, & Zagrovic, 2016), a collection of programs for the computation-intensive estimation of configurational entropy by information theoretical approaches on parallel architectures. All MD trajectories were first converted from Cartesian to Bond-Angle-Torsion (BAT) coordinates. To assess the convergence of configurational entropy (*S*_conf_), cumulative plots were generated for single copy systems using a 50-ns time step. Due to the relatively slow convergence of the entropy values (Figure S3B), the final entropy calculations were performed for the entire 1-µs trajectories. Note that the absolute *S*_conf_ values are negative and carry arbitrary units due to the exclusion from the calculations of the constant momentum part of the configurational entropy integrals and are reported just to illustrate the convergence of the entropy values as a function of simulated time. However, upon subtraction of these absolute values (i. e. for single and crowded systems), the relative entropy (Δ*S*_conf_) carries correct physical units and is equal to the total configurational entropy change between the two systems. The relative entropies of a protein in single-chain and multi-chain systems were averaged over all possible combinations of the two single-protein copies and the 24 crowded-system copies. To estimate the effect of mutations on configurational heterogeneity, the corresponding entropy differences were calculated as follows:

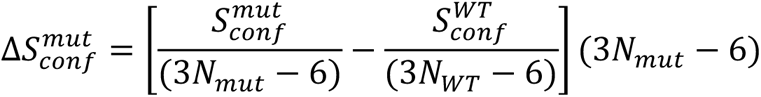

where *N*_*mut*_ and *N*_*WT*_ represent the numbers of atoms in mutant and WT proteins, respectively. The final values of configurational entropy differences were multiplied by the temperature (T = 310 K) and converted to kcal/mol units.

### Estimation of diffusion coefficients and shear viscosity

Diffusion coefficients of individual protein chains were calculated following the procedure described elsewhere (von Bülow, Siggel, Linke, & Hummer, 2019), together with viscosity estimation (Hess, 2002), application of corrections for size-dependent effects (Yeh & Hummer, 2004) and rescaling against experimentally comparable values (Fennell, Ghousifam, Haseleu, & Gappa-Fahlenkamp, 2018).

Thus, translational diffusion coefficients 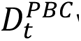 were extracted for individual molecules by analyzing center-of-mass mean-square displacement (MSD) curves, considering that:

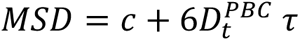

for τ approaching infinity. The above equation was fitted in a linear regime of MSD between 20 ns and 40 ns for the 24-copy systems (Figure S4D), and between 5 ns and 15 ns for the single molecule. As previously suggested (Yeh & Hummer, 2004), the thus obtained diffusion coefficients were corrected for size-dependent effects that arise from periodic boundary conditions. Applying this correction, the diffusion coefficient *D*_*t*_ can be determined as:

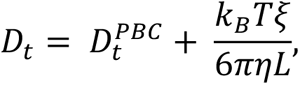

where L is the edge length of the simulation box, η is the viscosity of the system that the particle is simulated in, and ξ = 2.837297, a term arising from the cubic lattice (Yeh & Hummer, 2004). The latter correction requires estimation of shear viscosity values in the system. For this purpose, short 10 ns NVT MD simulation were performed for each system starting from the last snapshot of 1 µs simulations with detailed output for GROMACS energy-file (every 10 fs). Shear viscosities were extracted from these NVT simulations using the Green-Kubo formula (Hess, 2002):

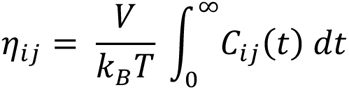

where V denotes the volume of the simulation box and *C*_i<_(*t*) is the autocorrelation function:

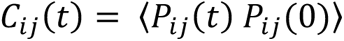

of the pressure tensor elements 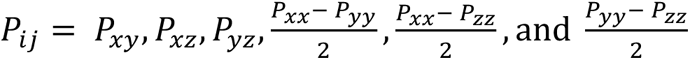

The autocorrelation function was numerically integrated between 0 ps and 1 ps, followed by analytical integration up to infinity. The analytical part of the integral was determined by a double exponential fit of the data between 1 ps and 5 ps (Figure S4E):

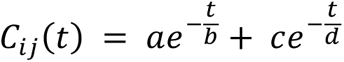

Shear viscosity η was then determined by averaging over the η_ij_ of the evaluated pressure tensor elements. Finally, the corrected diffusion coefficients were rescaled by the ratio of the simulated and experimentally determined water viscosities (Fennell et al., 2018):

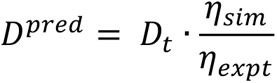

For the experimental water viscosity value at 310 K and 0.1 M salt *η*_*expt*_ of 0.69 mPa·s (Fennell et al., 2018) was used. The simulated value of viscosity (*η*_*sim*_= 0.83 mPa·s) in TIP4P-D water box at the same conditions was obtained previously by us (in preparation) using a series of 100 ns NVT MD simulations for cubic boxes of different size (3, 4, and 5 nm).

MSD curves for complete 1-µs MD trajectories or only their last 0.3 µs fragments were calculated using *msd* utility from the GROMACS package. Pressure tensors were obtained using *energy* utility from the GROMACS package for the analysis of NVT simulations. Viscosities were calculated as described above using MATLAB (R2009) scripts written specifically for this purpose.

### Modeling and visualization of condensates topology

To generate a visual representation of a condensate with a fractal dimension obtained by the self-propagation model (see Appendix), FracVAL algorithm was used (Morán et al., 2019). FracVAL is a tunable algorithm for generation of fractal structures of aggregates of polydisperse primary particles, which preserves the predefined fractal dimension (*d*_f_) and the fractal prefactor (*k*_f_) to generate aggregates of desired size. The scaling law in this case can be defined as shown in eq. 14. The prefactor *k*_4_ is equal to 1 in the present model (see Appendix). The *d*_4_ values for generating cluster models in FracVAL were calculated according to eq. 10 using the averaged *φ* and *n* values over the last 0.3 µs of the 24-copy MD simulations, while <*Rg_MD_>* values over the last 0.3 µs were taken as effective sizes of primary particles (detailed parameters are listed in Figure S4). For visualization purposes, the size of condensates generated by FracVAL was defined as 1024 molecules. FracVAL cluster models were transformed to all-atom resolution by using selected protein MD conformations with the respective *Rg* values and scripts specially written for this purpose. The obtained coarse-grained and all-atom structures were visualized using PyMol (Schrodinger & DeLano W, 2020).

## ACKNOWLEDGEMENTS

This work was supported by Austrian Science Fund FWF Standalone Grants P 30550 and P 30680-B21 (to B.Z.); a NOMIS Pioneering Research Grant and a grant of the Austrian Science Fund (FWF, project F79) (to A.K.).

## APPENDIX

### Fractal model of condensate assembly

The number of molecules in a cluster at iteration *i* can be calculated according to a simple geometrical progression:

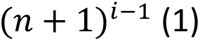

where *n* is the valency of interactions or more generally – the coordination number. An apparent volume *v* of a single molecule, whereby atoms occupy the fraction **φ** of that volume, can be expressed as:

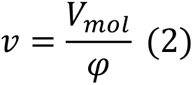

where *V*_*mol*_ is the molecular volume, which in turn is proportional to the molecular weight (*M*_W_) with a factor *k* (*k* = 1.21 used for all calculations(Harpaz, Gerstein, & Chothia, 1994)):

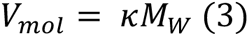

Thus, combination of (1) and (2) allows to estimate an apparent volume of condensate (*V*_i_) at iteration *i*:

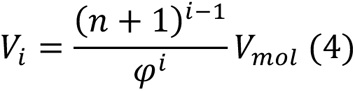

Under the assumption of a spherical geometry, a characteristic size of the condensate at iteration *i* (*R*_i_) can be derived from (4):

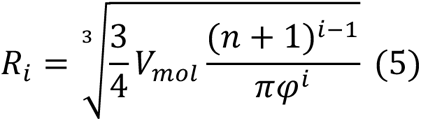

An effective molar concentration (*C*_i_) of molecules in a condensate at iteration *i* then can be estimated as:

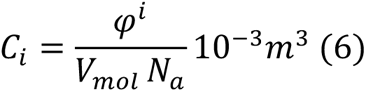

where, *N*_*a*_ is the Avogadro number. A combination of (5) and (6) gives the volume fraction as a function of valency for a condensate with a given concentration (*C*) and size (*R*):

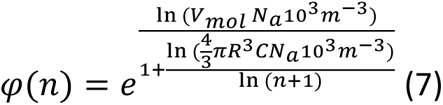

The above formalism allows one to extract the characteristic parameters **φ** and *n* from the linear regression of an empirical log *R* vs. log *M* plot. First, the mass *M*_*i*_ of a condensate at iteration *i* is given as:

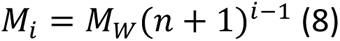

A combination of (5) and (8) then gives analytical expressions for a slope (*A*) and an intercept (*B*) for the linear regression plot:

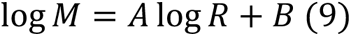

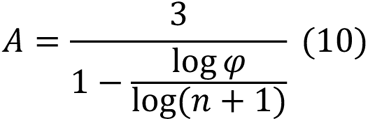

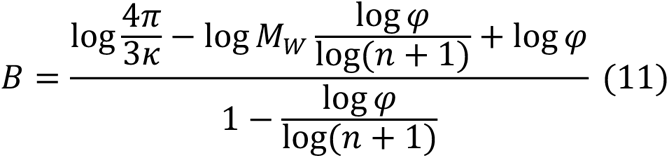

Finally, a combination of (10) and (11) gives equations for **φ** and *n* using the slope *A* and the intercept *B* for the linear regression (9):

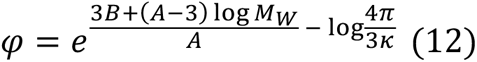

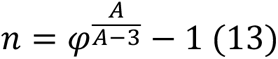

Note that *A*, also known as the “fractal dimension” (*d*_*f*_) (Carpineti & Giglio, 1992) is the key parameter describing the condensate organization. The scaling law in this case is typically defined as:

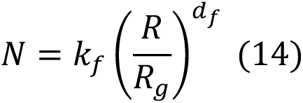

where *N* is the number of molecules in each cluster with the corresponding size *R*, while *R*_*g*_ is an effective size of an individual molecule, and *k*_*f*_is the fractal prefactor. In the present model of condensate assembly, *k*_*f*_ is equal to 1, as proven below. Thus, according to (14) the number of molecules for a cluster obtained at iteration *i* can be calculated as:

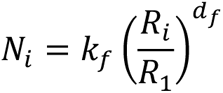

where *d*_4_ is a slope (*A*) for log *R* vs. log *M* linear regression and is defined in (10). A combination of (1) and (5) gives:

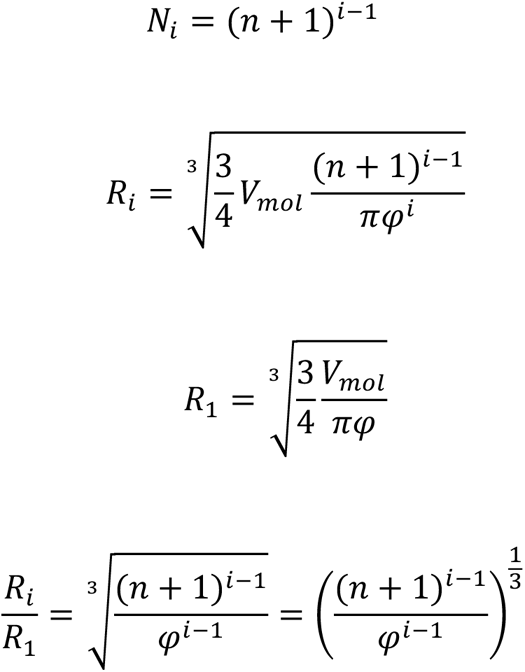

then simplification leads to:

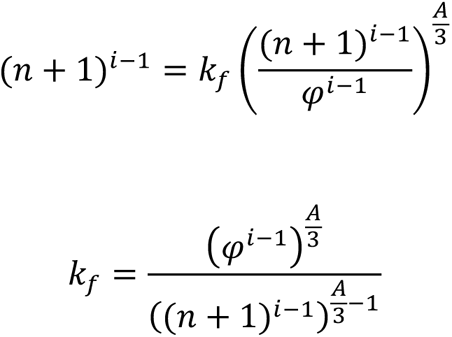

and by using (13) to substitute *n* + 1 one derives:

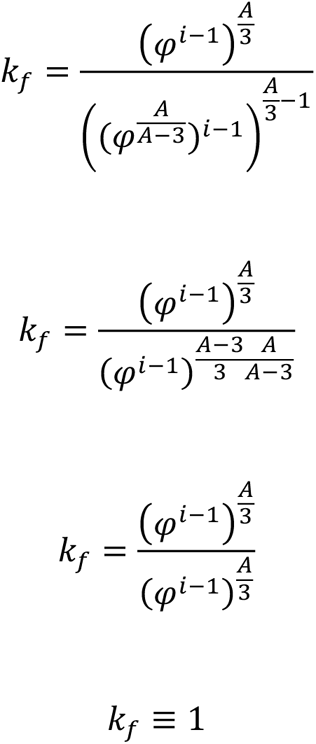

## REFERENCES

Abraham, M. J., Murtola, T., Schulz, R., Páll, S., Smith, J. C., Hess, B., & Lindahl, E. (2015). GROMACS: High performance molecular simulations through multi-level parallelism from laptops to supercomputers. SoftwareX, 1–2, 19-25. doi:10.1016/j.softx.2015.06.001

Alberti, S., & Dormann, D. (2019). Liquid-Liquid Phase Separation in Disease. Annu Rev Genet, 53, 171–194. doi:10.1146/annurev-genet-112618-043527

Alberti, S., & Hyman, A. A. (2021). Biomolecular condensates at the nexus of cellular stress, protein aggregation disease and ageing. Nature Reviews Molecular Cell Biology, 22(3), 196–213. doi:10.1038/s41580-020-00326-6

Altieri, A. S., Hinton, D. P., & Byrd, R. A. (1995). Association of Biomolecular Systems via Pulsed Field Gradient NMR Self-Diffusion Measurements. Journal of the American Chemical Society, 117(28), 7566–7567. doi:10.1021/ja00133a039

Armstrong, J. K., Wenby, R. B., Meiselman, H. J., & Fisher, T. C. (2004). The Hydrodynamic Radii of Macromolecules and Their Effect on Red Blood Cell Aggregation. Biophysical Journal, 87(6), 4259–4270. doi:10.1529/biophysj.104.047746

Banani, S. F., Lee, H. O., Hyman, A. A., & Rosen, M. K. (2017). Biomolecular condensates: organizers of cellular biochemistry. Nat Rev Mol Cell Biol, 18(5), 285–298. doi:10.1038/nrm.2017.7

Baum, M., Erdel, F., Wachsmuth, M., & Rippe, K. (2014). Retrieving the intracellular topology from multi-scale protein mobility mapping in living cells. Nature Communications, 5(1), 4494. doi:10.1038/ncomms5494

Benayad, Z., von Bülow, S., Stelzl, L. S., & Hummer, G. (2021). Simulation of FUS Protein Condensates with an Adapted Coarse-Grained Model. Journal of Chemical Theory and Computation, 17(1), 525–537. doi:10.1021/acs.jctc.0c01064

Bernado, P., & Blackledge, M. (2009). A self-consistent description of the conformational behavior of chemically denatured proteins from NMR and small angle scattering. Biophys J, 97(10), 2839–2845. doi:10.1016/j.bpj.2009.08.044

Boeynaems, S., Alberti, S., Fawzi, N. L., Mittag, T., Polymenidou, M., Rousseau, F., . . .Fuxreiter, M. (2018). Protein Phase Separation: A New Phase in Cell Biology. Trends Cell Biol, 28(6), 420–435. doi:10.1016/j.tcb.2018.02.004

Brady, J. P., Farber, P. J., Sekhar, A., Lin, Y.-H., Huang, R., Bah, A., . . . Kay, L. E. (2017). Structural and hydrodynamic properties of an intrinsically disordered region of a germ cell-specific protein on phase separation. Proceedings of the National Academy of Sciences, 114(39), E8194–E8203. doi:10.1073/pnas.1706197114

Brangwynne, C. P., Eckmann, C. R., Courson, D. S., Rybarska, A., Hoege, C., Gharakhani, J., . . . Hyman, A. A. (2009). Germline P Granules Are Liquid Droplets That Localize by Controlled Dissolution/Condensation. Science, 324(5935), 1729–1732. doi:10.1126/science.1172046

Brangwynne, C. P., Tompa, P., & Pappu, R. V. (2015). Polymer physics of intracellular phase transitions. Nature Physics, 11(11), 899–904. doi:10.1038/nphys3532

Bremer, A., Farag, M., Borcherds, W. M., Peran, I., Martin, E. W., Pappu, R. V., & Mittag, T. (2022). Deciphering how naturally occurring sequence features impact the phase behaviours of disordered prion-like domains. Nature Chemistry, 14(2), 196–207. doi:10.1038/s41557-021-00840-w

Carpineti, M., & Giglio, M. (1992). Spinodal-type dynamics in fractal aggregation of colloidal clusters. Phys Rev Lett, 68(22), 3327–3330. doi:10.1103/PhysRevLett.68.3327

Choi, J. M., Holehouse, A. S., & Pappu, R. V. (2020). Physical Principles Underlying the Complex Biology of Intracellular Phase Transitions. Annu Rev Biophys, 49, 107–133. doi:10.1146/annurev-biophys-121219-081629

Choi, J. M., Hyman, A. A., & Pappu, R. V. (2020). Generalized models for bond percolation transitions of associative polymers. Physical Review E, 102(4), 042403. doi:10.1103/PhysRevE.102.042403

Conicella, A. E., Dignon, G. L., Zerze, G. H., Schmidt, H. B., D’Ordine, A. M., Kim, Y. C., . . . Fawzi, N. L. (2020). TDP-43 alpha-helical structure tunes liquid-liquid phase separation and function. Proc Natl Acad Sci U S A, 117(11), 5883–5894. doi:10.1073/pnas.1912055117

da Silva, M. A., & Arêas, E. P. (2005). Solvent-induced lysozyme gels: rheology, fractal analysis, and sol-gel kinetics. J Colloid Interface Sci, 289(2), 394–401. doi:10.1016/j.jcis.2005.04.026

de Ruiter, A., Polyansky, A. A., & Zagrovic, B. (2017). Dependence of Binding Free Energies between RNA Nucleobases and Protein Side Chains on Local Dielectric Properties. J Chem Theory Comput, 13(9), 4504–4513. doi:10.1021/acs.jctc.6b01202

Dewar, M. J. S., Zoebisch, E. G., Healy, E. F., & Stewart, J. J. P. (1985). Development and use of quantum mechanical molecular models. 76. AM1: a new general purpose quantum mechanical molecular model. Journal of the American Chemical Society, 107(13), 3902–3909. doi:10.1021/ja00299a024

Dignon, G. L., Zheng, W., Best, R. B., Kim, Y. C., & Mittal, J. (2018). Relation between single-molecule properties and phase behavior of intrinsically disordered proteins. Proc Natl Acad Sci U S A, 115(40), 9929–9934. doi:10.1073/pnas.1804177115

Dror, R. O., Dirks, R. M., Grossman, J. P., Xu, H., & Shaw, D. E. (2012). Biomolecular Simulation: A Computational Microscope for Molecular Biology. Annual Review of Biophysics, 41(1), 429–452. doi:10.1146/annurev-biophys-042910-155245

Dubreuil, B., Matalon, O., & Levy, E. D. (2019). Protein Abundance Biases the Amino Acid Composition of Disordered Regions to Minimize Non-functional Interactions. Journal of Molecular Biology, 431(24), 4978–4992. doi:10.1016/j.jmb.2019.08.008

Dzuricky, M., Rogers, B. A., Shahid, A., Cremer, P. S., & Chilkoti, A. (2020). De novo engineering of intracellular condensates using artificial disordered proteins. Nature Chemistry, 12(9), 814–825. doi:10.1038/s41557-020-0511-7

Elbaum-Garfinkle, S., Kim, Y. C., Szczepaniak, K., Chen, C. C.-H., Eckmann, C. R., Myong, S., & Brangwynne, C. P. (2015). The disordered P granule protein LAF-1 drives phase separation into droplets with tunable viscosity and dynamics. Proceedings of the National Academy of Sciences, 112(23), 7189–7194. doi:10.1073/pnas.1504822112

Espinosa, J. R., Joseph, J. A., Sanchez-Burgos, I., Garaizar, A., Frenkel, D., & Collepardo-Guevara, R. (2020). Liquid network connectivity regulates the stability and composition of biomolecular condensates with many components. Proceedings of the National Academy of Sciences, 117(24), 13238–13247. doi:10.1073/pnas.1917569117

Feng, Z., Chen, X., Wu, X., & Zhang, M. (2019). Formation of biological condensates via phase separation: Characteristics, analytical methods, and physiological implications. J Biol Chem, 294(40), 14823–14835. doi:10.1074/jbc.REV119.007895

Fennell, C. J., Ghousifam, N., Haseleu, J. M., & Gappa-Fahlenkamp, H. (2018). Computational Signaling Protein Dynamics and Geometric Mass Relations in Biomolecular Diffusion. J Phys Chem B, 122(21), 5599–5609. doi:10.1021/acs.jpcb.7b11846

Fisher, R. S., & Elbaum-Garfinkle, S. (2020). Tunable multiphase dynamics of arginine and lysine liquid condensates. Nature Communications, 11(1), 4628. doi:10.1038/s41467-020-18224-y

Fleck, M., Polyansky, A. A., & Zagrovic, B. (2016). PARENT: A Parallel Software Suite for the Calculation of Configurational Entropy in Biomolecular Systems. Journal of Chemical Theory and Computation, 12(4), 2055–2065. doi:10.1021/acs.jctc.5b01217

Fleck, M., Polyansky, A. A., & Zagrovic, B. (2018). Self-Consistent Framework Connecting Experimental Proxies of Protein Dynamics with Configurational Entropy. Journal of Chemical Theory and Computation, 14(7), 3796–3810. doi:10.1021/acs.jctc.8b00100

Forrest, S. R., & Witten, T. A. (1979). Long-range correlations in smoke-particle aggregates. Journal of Physics A: Mathematical and General, 12(5), L109–L117. doi:10.1088/0305-4470/12/5/008

Fossat, M. J., Zeng, X., & Pappu, R. V. (2021). Uncovering Differences in Hydration Free Energies and Structures for Model Compound Mimics of Charged Side Chains of Amino Acids. The Journal of Physical Chemistry B, 125(16), 4148–4161. doi:10.1021/acs.jpcb.1c01073

Gallego, L. D., Schneider, M., Mittal, C., Romanauska, A., Gudino Carrillo, R. M., Schubert, T., . . . Kohler, A. (2020). Phase separation directs ubiquitination of gene-body nucleosomes. Nature, 579(7800), 592–597. doi:10.1038/s41586-020-2097-z

Garcia-Jove, N. M., Kashida, S., Chouaib, R., Souquere, S., Pierron, G., Weil, D., & Gueroui, Z. (2019). RNA is a critical element for the sizing and the composition of phase-separated RNA–protein condensates. Nature Communications, 10(1), 3230. doi:10.1038/s41467-019-11241-6

Gonçalves, A. D., Alexander, C., Roberts, C. J., Spain, S. G., Uddin, S., & Allen, S. (2016). The effect of protein concentration on the viscosity of a recombinant albumin solution formulation. RSC Advances, 6(18), 15143–15154. doi:10.1039/C5RA21068B

Guillén-Boixet, J., Kopach, A., Holehouse, A. S., Wittmann, S., Jahnel, M., Schlüßler, R., . . . Franzmann, T. M. (2020). RNA-Induced Conformational Switching and Clustering of G3BP Drive Stress Granule Assembly by Condensation. Cell, 181(2), 346–361.e317. doi:10.1016/j.cell.2020.03.049

Hagiwara, T., Kumagai, H., & Nakamura, K. (1996). Fractal Analysis of Aggregates Formed by Heating Dilute BSA Solutions Using Light Scattering Methods. *Bioscience*, Biotechnology, and Biochemistry, 60(11), 1757–1763. doi:10.1271/bbb.60.1757

Harpaz, Y., Gerstein, M., & Chothia, C. (1994). Volume changes on protein folding. Structure, 2(7), 641–649. doi:10.1016/s0969-2126(00)00065-4

Havlin, S., & Ben-Avraham, D. (1987). Diffusion in disordered media. Advances in Physics, 36(6), 695–798. doi:10.1080/00018738700101072

Hess, B. (2002). Determining the shear viscosity of model liquids from molecular dynamics simulations. The Journal of Chemical Physics, 116(1), 209–217. doi:10.1063/1.1421362

Hess, B., Bekker, H., Berendsen, H. J. C., & Fraaije, J. G. E. M. (1997). LINCS: A linear constraint solver for molecular simulations. Journal of Computational Chemistry, 18(12), 1463–1472. doi:Doi 10.1002/(Sici)1096-987x(199709)18:12<1463::Aid-Jcc4>3.0.Co;2-H

Hong, Y., Najafi, S., Casey, T., Shea, J.-E., Han, S.-I., & Hwang, D. S. (2022). Hydrophobicity of arginine leads to reentrant liquid-liquid phase separation behaviors of arginine-rich proteins. Nature Communications, 13(1), 7326. doi:10.1038/s41467-022-35001-1

Hoover, W. G. (1985). Canonical dynamics: Equilibrium phase-space distributions. Phys Rev A Gen Phys, 31(3), 1695–1697. Retrieved from http://www.ncbi.nlm.nih.gov/pubmed/9895674

Jain, A., & Vale, R. D. (2017). RNA phase transitions in repeat expansion disorders. Nature, 546(7657), 243–247. doi:10.1038/nature22386

Jorgensen, W. L., Chandrasekhar, J., Madura, J. D., Impey, R. W., & Klein, M. L. (1983). Comparison of simple potential functions for simulating liquid water. The Journal of Chemical Physics, 79(2), 926–935. doi:10.1063/1.445869

Jorgensen, W. L., Maxwell, D. S., & Tirado-Rives, J. (1996). Development and Testing of the OPLS All-Atom Force Field on Conformational Energetics and Properties of Organic Liquids. Journal of the American Chemical Society, 118(45), 11225–11236. doi:10.1021/ja9621760

Jorgensen, W. L., & Ravimohan, C. (1985). Monte Carlo simulation of differences in free energies of hydration. The Journal of Chemical Physics, 83(6), 3050–3054. doi:10.1063/1.449208

Jorgensen, W. L., & Tirado-Rives, J. (2005). Molecular modeling of organic and biomolecular systems using BOSS and MCPRO. Journal of Computational Chemistry, 26(16), 1689–1700. doi:10.1002/jcc.20297

Kapitulnik, A., Gefen, Y., & Aharony, A. (1984). On the Fractal dimension and correlations in percolation theory. Journal of Statistical Physics, 36(5), 807–814. doi:10.1007/BF01012940

Kar, M., Dar, F., Welsh, T. J., Vogel, L. T., Kühnemuth, R., Majumdar, A., . . . Pappu, R. V. (2022). Phase-separating RNA-binding proteins form heterogeneous distributions of clusters in subsaturated solutions. Proceedings of the National Academy of Sciences, 119(28), e2202222119. doi:10.1073/pnas.2202222119

Kätzel, U., Bedrich, R., Stintz, M., Ketzmerick, R., Gottschalk-Gaudig, T., & Barthel, H. (2008). Dynamic Light Scattering for the Characterization of Polydisperse Fractal Systems: I. Simulation of the Diffusional Behavior. Particle & Particle Systems Characterization, 25, 9–18.

Keating, C. D., & Pappu, R. V. (2021). Liquid–Liquid Phase Separation: A Widespread and Versatile Way to Organize Aqueous Solutions. The Journal of Physical Chemistry Letters, 12(45), 10994–10995. doi:10.1021/acs.jpclett.1c03352

Kelley, L. A., Mezulis, S., Yates, C. M., Wass, M. N., & Sternberg, M. J. (2015). The Phyre2 web portal for protein modeling, prediction and analysis. Nat Protoc, 10(6), 845–858. doi:10.1038/nprot.2015.053

Kharlamova, A., Nicolai, T., & Chassenieux, C. (2020). Gelation of whey protein fractal aggregates induced by the interplay between added HCl, CaCl2 and NaCl. International Dairy Journal, *111*, 104824. doi:10.1016/j.idairyj.2020.104824

Khatun, S., Singh, A., Maji, S., Maiti, T. K., Pawar, N., & Gupta, A. N. (2020). Fractal self-assembly and aggregation of human amylin. Soft Matter, 16(12), 3143–3153. doi:10.1039/C9SM02463H

King, B. M., Silver, N. W., & Tidor, B. (2012). Efficient Calculation of Molecular Configurational Entropies Using an Information Theoretic Approximation. The Journal of Physical Chemistry B, 116(9), 2891–2904. doi:10.1021/jp2068123

King, J. T., & Shakya, A. (2021). Phase separation of DNA: From past to present. Biophysical Journal, 120(7), 1139–1149. doi:https://doi.org/10.1016/j.bpj.2021.01.033

Klein, R., Weitz, D. A., Lin, M. Y., Lindsay, H. M., Ball, R. C., & Meakin, P. (1990, 1990//). Theory of scattering from colloidal aggregates. Paper presented at the Trends in Colloid and Interface Science IV, Darmstadt.

Knowles, T. P., Fitzpatrick, A. W., Meehan, S., Mott, H. R., Vendruscolo, M., Dobson, C. M., & Welland, M. E. (2007). Role of intermolecular forces in defining material properties of protein nanofibrils. Science, 318(5858), 1900–1903. doi:10.1126/science.1150057

Krainer, G., Welsh, T. J., Joseph, J. A., Espinosa, J. R., Wittmann, S., de Csilléry, E., . . . Knowles, T. P. J. (2021). Reentrant liquid condensate phase of proteins is stabilized by hydrophobic and non-ionic interactions. Nature Communications, 12(1), 1085. doi:10.1038/s41467-021-21181-9

Lafontaine, D. L. J., Riback, J. A., Bascetin, R., & Brangwynne, C. P. (2021). The nucleolus as a multiphase liquid condensate. Nature Reviews Molecular Cell Biology, 22(3), 165–182. doi:10.1038/s41580-020-0272-6

Lasker, K., Boeynaems, S., Lam, V., Stainton, E., Jacquemyn, M., Daelemans, D., . . . Shapiro, L. (2021). A modular platform for engineering function of natural and synthetic biomolecular condensates. bioRxiv, 2021.2002.2003.429226. doi:10.1101/2021.02.03.429226

Lazzari, S., Nicoud, L., Jaquet, B., Lattuada, M., & Morbidelli, M. (2016). Fractal-like structures in colloid science. Adv Colloid Interface Sci, 235, 1–13. doi:10.1016/j.cis.2016.05.002

Lewis, R. N. A. H., & McElhaney, R. N. (2013). Membrane lipid phase transitions and phase organization studied by Fourier transform infrared spectroscopy. Biochimica et Biophysica Acta (BBA) - Biomembranes, 1828(10), 2347–2358. doi:10.1016/j.bbamem.2012.10.018

Li, L., Casalini, T., Arosio, P., & Salvalaglio, M. (2022). Modeling the Structure and Interactions of Intrinsically Disordered Peptides with Multiple Replica, Metadynamics-Based Sampling Methods and Force-Field Combinations. Journal of Chemical Theory and Computation, 18(3), 1915–1928. doi:10.1021/acs.jctc.1c00889

Lin, M. Y., Lindsay, H. M., Weitz, D. A., Ball, R. C., Klein, R., & Meakin, P. (1989). Universality in colloid aggregation. Nature, 339(6223), 360–362. doi:10.1038/339360a0

Lin, Y., Protter, D. S., Rosen, M. K., & Parker, R. (2015). Formation and Maturation of Phase-Separated Liquid Droplets by RNA-Binding Proteins. Molecular cell, 60(2), 208–219. doi:10.1016/j.molcel.2015.08.018

Lindorff-Larsen, K., Piana, S., Palmo, K., Maragakis, P., Klepeis, J. L., Dror, R. O., & Shaw, D. E. (2010). Improved side-chain torsion potentials for the Amber ff99SB protein force field. Proteins, 78(8), 1950–1958. doi:10.1002/prot.22711

Martin, E. W., Holehouse, A. S., Peran, I., Farag, M., Incicco, J. J., Bremer, A., . . . Mittag, T. (2020). Valence and patterning of aromatic residues determine the phase behavior of prion-like domains. Science, 367(6478), 694–699. doi:10.1126/science.aaw8653

McCall, P. M., Kim, K., Fritsch, A. W., Iglesias-Artola, J. M., Jawerth, L. M., Wang, J., . . . Brugués, J. (2020). Quantitative phase microscopy enables precise and efficient determination of biomolecular condensate composition. *bioRxiv*, 2020.2010.2025.352823. doi:10.1101/2020.10.25.352823

McSwiggen, D. T., Mir, M., Darzacq, X., & Tjian, R. (2019). Evaluating phase separation in live cells: diagnosis, caveats, and functional consequences. Genes & development, 33(23-24), 1619–1634. doi:10.1101/gad.331520.119

Mitrea, D. M., & Kriwacki, R. W. (2016). Phase separation in biology; functional organization of a higher order. Cell Communication and Signaling, 14(1), 1. doi:10.1186/s12964-015-0125-7

Mittag, T., & Pappu, R. V. (2022). A conceptual framework for understanding phase separation and addressing open questions and challenges. Molecular cell, 82(12), 2201–2214. doi:10.1016/j.molcel.2022.05.018

Molliex, A., Temirov, J., Lee, J., Coughlin, M., Kanagaraj, A. P., Kim, H. J., . . . Taylor, J. P. (2015). Phase separation by low complexity domains promotes stress granule assembly and drives pathological fibrillization. Cell, 163(1), 123–133. doi:10.1016/j.cell.2015.09.015

Morán, J., Fuentes, A., Liu, F., & Yon, J. (2019). FracVAL: An improved tunable algorithm of cluster–cluster aggregation for generation of fractal structures formed by polydisperse primary particles. Computer Physics Communications, 239, 225–237. doi:10.1016/j.cpc.2019.01.015

Murthy, A. C., Dignon, G. L., Kan, Y., Zerze, G. H., Parekh, S. H., Mittal, J., & Fawzi, N. L. (2019). Molecular interactions underlying liquid-liquid phase separation of the FUS low-complexity domain. Nat Struct Mol Biol, 26(7), 637–648. doi:10.1038/s41594-019-0250-x

Musacchio, A. (2022). On the role of phase separation in the biogenesis of membraneless compartments. The EMBO Journal, 41(5), e109952. doi:10.15252/embj.2021109952

Nicoud, L., Lazzari, S., Balderas Barragán, D., & Morbidelli, M. (2015). Fragmentation of amyloid fibrils occurs in preferential positions depending on the environmental conditions. J Phys Chem B, 119(13), 4644–4652. doi:10.1021/acs.jpcb.5b01160

Nott, T. J., Petsalaki, E., Farber, P., Jervis, D., Fussner, E., Plochowietz, A., . . . Baldwin, A. J. (2015). Phase transition of a disordered nuage protein generates environmentally responsive membraneless organelles. Molecular cell, 57(5), 936–947. doi:10.1016/j.molcel.2015.01.013

Paloni, M., Bailly, R., Ciandrini, L., & Barducci, A. (2020). Unraveling Molecular Interactions in Liquid-Liquid Phase Separation of Disordered Proteins by Atomistic Simulations. J Phys Chem B, 124(41), 9009–9016. doi:10.1021/acs.jpcb.0c06288

Paloni, M., Bussi, G., & Barducci, A. (2021). Arginine multivalency stabilizes protein/RNA condensates. Protein Science, 30(7), 1418–1426. doi:10.1002/pro.4109

Pappu, R. V., Cohen, S. R., Dar, F., Farag, M., & Kar, M. (2023). Phase Transitions of Associative Biomacromolecules. Chemical Reviews. doi:10.1021/acs.chemrev.2c00814

Parrinello, M., & Rahman, A. (1981). Polymorphic Transitions in Single-Crystals - a New Molecular-Dynamics Method. Journal of Applied Physics, 52(12), 7182–7190. doi:10.1063/1.328693

Piana, S., Donchev, A. G., Robustelli, P., & Shaw, D. E. (2015). Water dispersion interactions strongly influence simulated structural properties of disordered protein states. J Phys Chem B, 119(16), 5113–5123. doi:10.1021/jp508971m

Polyansky, A. A., Volynsky, P. E., Arseniev, A. S., & Efremov, R. G. (2009). Adaptation of a membrane-active peptide to heterogeneous environment. I. Structural plasticity of the peptide. J Phys Chem B, 113(4), 1107–1119. doi:10.1021/jp803640e

Rauscher, S., & Pomes, R. (2017). The liquid structure of elastin. Elife, 6. doi:10.7554/eLife.26526

Roberts, S., Harmon, T. S., Schaal, J. L., Miao, V., Li, K., Hunt, A., . . . Chilkoti, A. (2018). Injectable tissue integrating networks from recombinant polypeptides with tunable order. Nature Materials, 17(12), 1154–1163. doi:10.1038/s41563-018-0182-6

Ruff, K. M., Khan, S. J., & Pappu, R. V. (2014). A Coarse-Grained Model for Polyglutamine Aggregation Modulated by Amphipathic Flanking Sequences. Biophysical Journal, 107(5), 1226–1235. doi:10.1016/j.bpj.2014.07.019

Ryan, V. H., Dignon, G. L., Zerze, G. H., Chabata, C. V., Silva, R., Conicella, A. E., . . . Fawzi, N. L. (2018). Mechanistic View of hnRNPA2 Low-Complexity Domain Structure, Interactions, and Phase Separation Altered by Mutation and Arginine Methylation. Molecular cell, 69(3), 465–479 e467. doi:10.1016/j.molcel.2017.12.022

Sanders, D. W., Kedersha, N., Lee, D. S. W., Strom, A. R., Drake, V., Riback, J. A., . . . Brangwynne, C. P. (2020). Competing Protein-RNA Interaction Networks Control Multiphase Intracellular Organization. Cell, 181(2), 306–324.e328. doi:10.1016/j.cell.2020.03.050

Schmit, J. D., Bouchard, J. J., Martin, E. W., & Mittag, T. (2020). Protein Network Structure Enables Switching between Liquid and Gel States. Journal of the American Chemical Society, 142(2), 874–883. doi:10.1021/jacs.9b10066

Schrodinger, L., & DeLano W. (2020). PyMOL. Retrieved from http://www.pymol.org/pymol

Schuster, B. S., Dignon, G. L., Tang, W. S., Kelley, F. M., Ranganath, A. K., Jahnke, C. N., . . . Mittal, J. (2020). Identifying sequence perturbations to an intrinsically disordered protein that determine its phase-separation behavior. Proceedings of the National Academy of Sciences, 117(21), 11421–11431. doi:10.1073/pnas.2000223117

Seim, I., Posey, A. E., Snead, W. T., Stormo, B. M., Klotsa, D., Pappu, R. V., & Gladfelter, A. S. (2022). Dilute phase oligomerization can oppose phase separation and modulate material properties of a ribonucleoprotein condensate. Proc Natl Acad Sci U S A, 119(13), e2120799119. doi:10.1073/pnas.2120799119

Slomkowski, S., Alemán, J. V., Gilbert, R. G., Hess, M., Horie, K., Jones, R. G., . . . Stepto, R. F. T. (2011). Terminology of polymers and polymerization processes in dispersed systems (IUPAC Recommendations 2011). Pure and Applied Chemistry, 83(12), 2229–2259. doi:10.1351/PAC-REC-10-06-03

Sprague, B. L., & McNally, J. G. (2005). FRAP analysis of binding: proper and fitting. Trends Cell Biol, 15(2), 84–91. doi:10.1016/j.tcb.2004.12.001

Stanley, E. H. (1984). Application of fractal concepts to polymer statistics and to anomalous transport in randomly porous media. Journal of Statistical Physics, 36(5), 843–860. doi:10.1007/BF01012944

Stauffer, D., Coniglio, A., & Adam, M. (1982, 1982//). *Gelation and critical phenomena.* Paper presented at the Polymer Networks, Berlin, Heidelberg.

Taylor, N. O., Wei, M. T., Stone, H. A., & Brangwynne, C. P. (2019). Quantifying Dynamics in Phase-Separated Condensates Using Fluorescence Recovery after Photobleaching. Biophysical Journal, 117(7), 1285–1300. doi:10.1016/j.bpj.2019.08.030

Thouy, R., & Jullien, R. (1994). A cluster-cluster aggregation model with tunable fractal dimension. Journal of Physics A: Mathematical and General, 27(9), 2953–2963. doi:10.1088/0305-4470/27/9/012

Uversky, V. N. (2009). Intrinsically disordered proteins and their environment: effects of strong denaturants, temperature, pH, counter ions, membranes, binding partners, osmolytes, and macromolecular crowding. Protein J, 28(7-8), 305–325. doi:10.1007/s10930-009-9201-4

Uversky, V. N. (2021). Recent Developments in the Field of Intrinsically Disordered Proteins: Intrinsic Disorder–Based Emergence in Cellular Biology in Light of the Physiological and Pathological Liquid–Liquid Phase Transitions. Annual Review of Biophysics, 50(1), 135–156. doi:10.1146/annurev-biophys-062920-063704

Vernon, R. M., Chong, P. A., Tsang, B., Kim, T. H., Bah, A., Farber, P., . . . Forman-Kay, J. D. (2018). Pi-Pi contacts are an overlooked protein feature relevant to phase separation. Elife, 7. doi:10.7554/eLife.31486

von Bülow, S., Siggel, M., Linke, M., & Hummer, G. (2019). Dynamic cluster formation determines viscosity and diffusion in dense protein solutions. Proceedings of the National Academy of Sciences, 116(20), 9843–9852. doi:10.1073/pnas.1817564116

Wang, J., Choi, J. M., Holehouse, A. S., Lee, H. O., Zhang, X., Jahnel, M., . . . Hyman, A. A. (2018). A Molecular Grammar Governing the Driving Forces for Phase Separation of Prion-like RNA Binding Proteins. Cell, 174(3), 688–699 e616. doi:10.1016/j.cell.2018.06.006

Wei, M. T., Elbaum-Garfinkle, S., Holehouse, A. S., Chen, C. C., Feric, M., Arnold, C. B., . . . Brangwynne, C. P. (2017). Phase behaviour of disordered proteins underlying low density and high permeability of liquid organelles. Nat Chem, 9(11), 1118–1125. doi:10.1038/nchem.2803

Yeh, I.-C., & Hummer, G. (2004). System-Size Dependence of Diffusion Coefficients and Viscosities from Molecular Dynamics Simulations with Periodic Boundary Conditions. The Journal of Physical Chemistry B, 108(40), 15873–15879. doi:10.1021/jp0477147

Zaslavsky, B. Y., & Uversky, V. N. (2018). In Aqua Veritas: The Indispensable yet Mostly Ignored Role of Water in Phase Separation and Membrane-less Organelles. Biochemistry, 57(17), 2437–2451. doi:10.1021/acs.biochem.7b01215

Zeng, X., Ruff, K. M., & Pappu, R. V. (2022). Competing interactions give rise to two-state behavior and switch-like transitions in charge-rich intrinsically disordered proteins. Proceedings of the National Academy of Sciences, 119(19), e2200559119. doi:10.1073/pnas.2200559119

Zheng, W., Dignon, G. L., Jovic, N., Xu, X., Regy, R. M., Fawzi, N. L., . . . Mittal, J. (2020). Molecular Details of Protein Condensates Probed by Microsecond Long Atomistic Simulations. J Phys Chem B, 124(51), 11671–11679. doi:10.1021/acs.jpcb.0c10489

