## Supplementary material for "Protein compactness and interaction valency define the architecture of a biomolecular condensate across scales": Figure S1

**A**

| Name | Protein | Water | Na+ | Cl- | [Protein] (mM) | [Protein] (g/l) | Box size (nm) | MD time (μs) | Replicas |
|---|---|---|---|---|---|---|---|---|---|
| Lge1 1-80 WT | 1 | 23674 | 44 | 50 | 2.3 | 20.7 | 9.0 | 1 | 2 |
| Lge1 1-80 Y>A | 1 | 23742 | 44 | 50 | 2.3 | 17.7 | 9.0 | 1 | 2 |
| Lge1 1-80 R>K | 1 | 23673 | 44 | 50 | 2.3 | 20.0 | 9.0 | 1 | 2 |
| Lge1 1-80 24 copies | 24 | 182056 | 351 | 495 | 6.9 | 62.5 | 18.0 | 1 | 1 |
| Lge1 1-80 Y>A 24 copies | 24 | 183407 | 351 | 495 | 6.9 | 45.0 | 18.0 | 1 | 1 |
| Lge1 1-80 R>K 24 copies | 24 | 217857 | 413 | 557 | 5.8 | 50.8 | 19.0 | 1 | 1 |

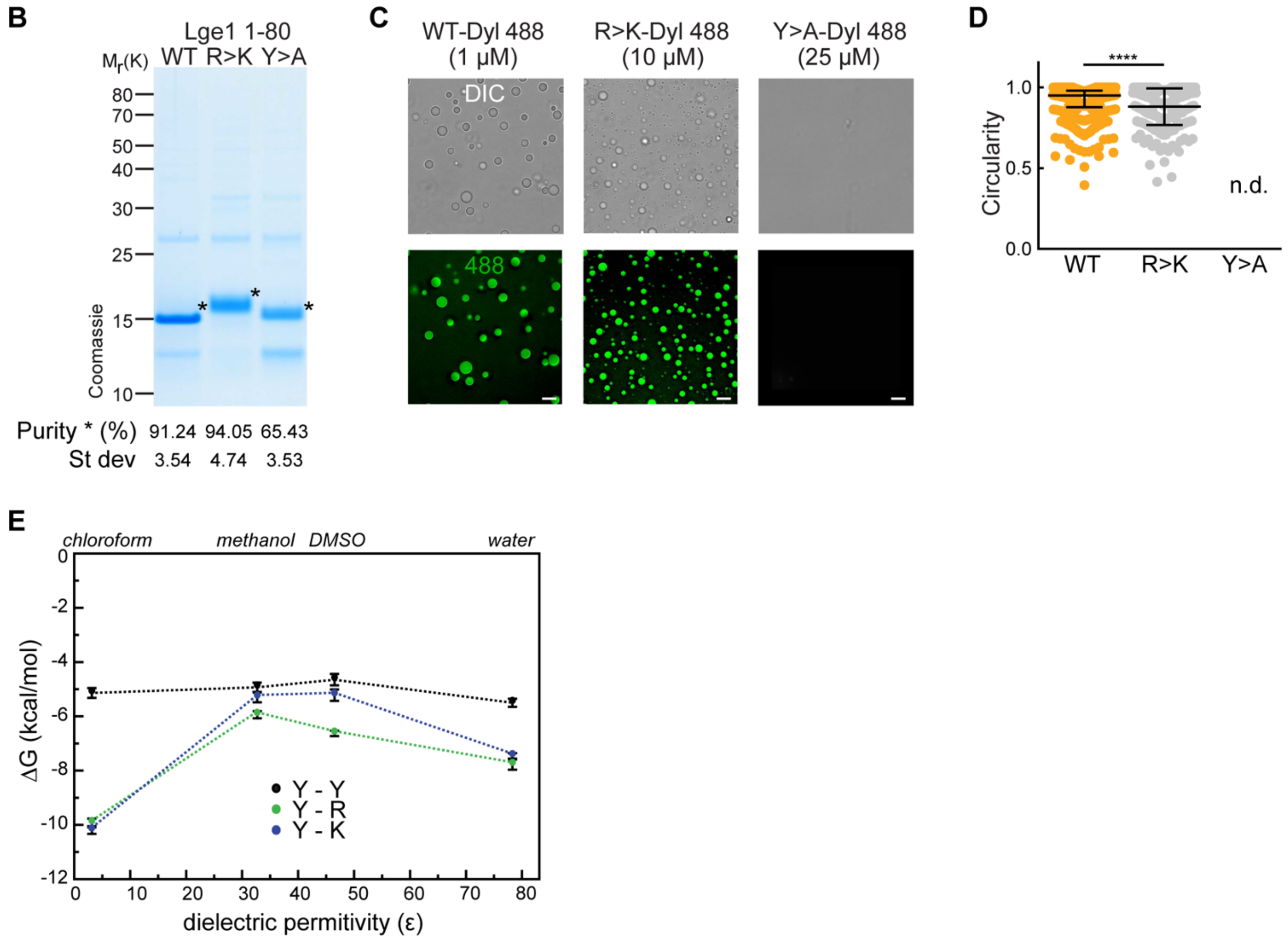


**Figure S1. Experimental and modeling studies of the effect of tyrosine and arginine mutations in Lge1$_{1-80}$.** (**A**). Details of simulated systems including composition, effective molar and mass protein concentration, size of the simulated cubic box, simulation time and the number of replicas. (**B**) Lge1$_{1-80}$ purification and purity. Following purification, the constructs were analyzed by SDS-Page (4-12 % gel, MOPS buffer) and Coomassie staining. Lge1$_{1-80}$ purity was assessed by densitometry, comparing Lge1 (*) to other impurities detected in the same lane (see Methods for the protocol). Average and standard deviation from 3 different purifications are included. (**C**) Representative images of Dylight-labelled Lge1$_{1-80}$ WT, R>K and Y>A, generated at the indicated protein concentrations. Scale bar, 5 μm. (**D**) Circularity of Dylight-labelled Lge1$_{1-80}$ WT (1 μM; *orange*) and R>K (10 μM; *light grey*) condensates in solution (25 mM Tris, 100 mM NaCl, pH 7.5). Median and interquartile range are indicated. $n_{WT}$ = 300, $n_{R>K}$ = 286. n.d., not determinable. ****p<0.0001, determined by two-sided Mann-Whitney test. (**E**) Dependence of the pairwise interaction free energies (*ΔG*) of arginine (R), lysine (K) and tyrosine (Y) side-chain analogs on the dielectric permittivity of the environment (*Ɛ*). Results of all-atom Monte-Carlo simulations (see Methods for more details).

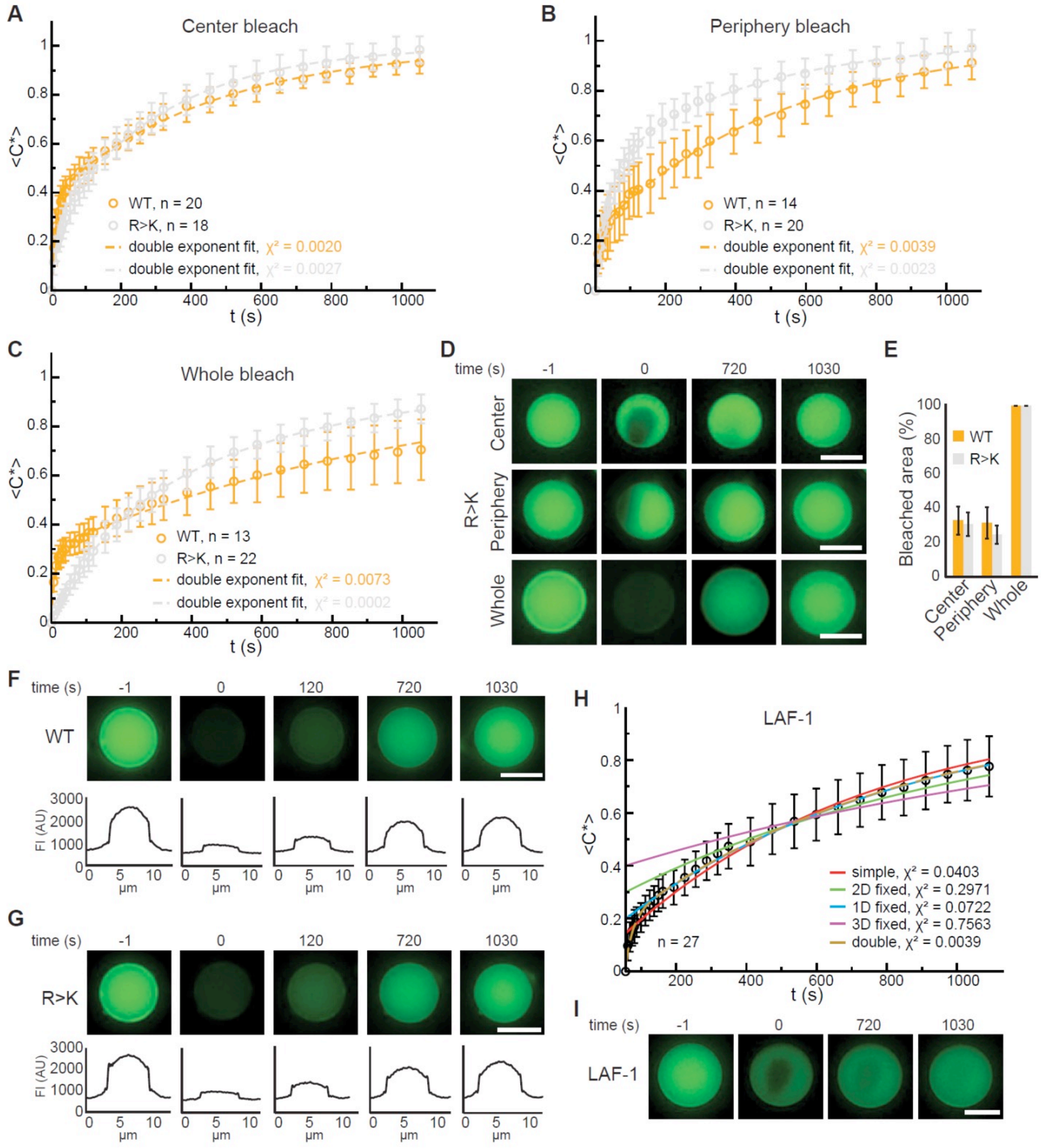


**Figure S2. FRAP analyses of Lge1$_{1-80}$ condensates**. FRAP curve of Dylight-labelled Lge1$_{1-80}$ WT (orange) and R>K (grey) condensates that were bleached in the center (**A**), the periphery (**B**) or across the whole condensate (**C**). Recovery of the normalized intensity inside the bleach spot <*C**> over time is shown as average (open circles) and standard deviation. The data is fitted to a double exponential model (dashed line). n, number of individual condensates. χ2, chi-squared value. (**D**) Representative FRAP images of Lge1$_{1-80}$ R>K condensates, bleached in the center (*upper panels*), the periphery (*middle*) or across the whole condensate (*lower panels*), including pre-bleach (*left, time -1 s*), bleach (*time 0 s*) and post-bleach (*time 720 s, 1030 s*). Scale bars, 5 µm. (**E**) Average of the percentage of the bleached area calculated over the total area of Lge1$_{1-80}$ WT (orange) and R>K (grey) condensates that were bleached in the center, periphery and across the whole condensate. Error bars represent standard deviation. Number of condensates quantified as in A, B, C. (**F**) Representative FRAP images of a Lge1$_{1-80}$ WT condensate that was bleached across the whole condensate, including pre-bleach (*time -1s*), bleach (*time 0s*) and post-bleach (*time 120 s, 720 s, 1030 s*). The corresponding line profiles of the condensate at different times after bleaching show homogeneous fluorescent recovery. FI, fluorescent intensity. Scale bar, 5 µm. (**G**) Representative FRAP images of a Lge1$_{1-80}$ R>K condensate that was bleached across the whole condensate. The fluorescent line profile shows the same recovery behavior as for Lge1$_{1-80}$ WT in F. FI, fluorescent intensity. Scale bar, 5 µm. (**H**) FRAP curve of Dylight-labelled LAF-1 shown as the average of the normalized intensity inside the bleach spot *<C*>* over time (black open circles). Error bars represent standard deviation. Fits to different models are plotted, with the corresponding chi-squared values (χ2). (**I**) Representative FRAP images of a LAF-1 condensate that was bleached in the center, including pre-bleach (*time -1s*), bleach (*time 0s*) and post-bleach (*time 720 s, 1030 s*). Scale bar, 5 µm.

**A** *crowded*

WT

| | % 24 | % 1 | Enr 24 | Enr 1 |
|---|---|---|---|---|
| R-Y | 10 | 6 | 2,1 | 1,2 |
| G-Y | 7 | 5 | 1,2 | 0,9 |
| Y-Y | 7 | 6 | 2,2 | 2,0 |
| S-Y | 4 | 4 | 1,2 | 1,2 |
| P-Y | 4 | 1 | 1,8 | 0,6 |
| T-Y | 4 | 2 | 1,7 | 0,9 |
| A-Y | 3 | 2 | 1,5 | 0,9 |
| N-Y | 3 | 3 | 1,4 | 1,2 |
| Q-Y | 3 | 1 | 1,5 | 0,8 |
| P-R | 2 | 2 | 1,4 | 1,0 |

R>K

| | % 24 | % 1 | Enr 24 | Enr 1 |
|---|---|---|---|---|
| G-Y | 11 | 6 | 1,8 | 1,0 |
| Y-Y | 10 | 7 | 3,1 | 2,3 |
| K-Y | 7 | 5 | 1,5 | 1,0 |
| T-Y | 5 | 2 | 2,4 | 0,9 |
| S-Y | 5 | 5 | 1,5 | 1,4 |
| A-Y | 4 | 3 | 2,0 | 1,2 |
| N-Y | 4 | 3 | 2,0 | 1,5 |
| P-Y | 4 | 2 | 1,9 | 0,7 |
| Q-Y | 3 | 2 | 1,7 | 0,9 |
| D-Y | 2 | 1 | 1,4 | 0,7 |

Y>A

| | % 24 | % 1 | Enr 24 | Enr 1 |
|---|---|---|---|---|
| A-R | 8 | 6 | 1,2 | 1,0 |
| A-T | 4 | 2 | 1,4 | 0,8 |
| A-Q | 3 | 1 | 1,3 | 0,6 |
| R-T | 3 | 2 | 1,8 | 1,1 |
| D-R | 3 | 1 | 2,9 | 1,2 |
| N-R | 3 | 2 | 1,7 | 1,0 |
| Q-R | 3 | 2 | 2,1 | 1,5 |
| R-S | 3 | 3 | 1,0 | 1,1 |
| G-T | 3 | 2 | 1,3 | 0,8 |
| R-R | 2 | 4 | 1,2 | 2,2 |

Enrichment of contacts over sequence background: 3,0 – 0,5

**B** *single*

WT

| | % 1 | % 24 | Enr 1 | Enr 24 |
|---|---|---|---|---|
| Y-Y | 6 | 7 | 2,0 | 2,2 |
| R-Y | 6 | 10 | 1,2 | 2,1 |
| G-G | 5 | 1 | 1,7 | 0,3 |
| S-Y | 4 | 4 | 1,2 | 1,2 |
| R-R | 4 | 1 | 1,9 | 0,7 |
| N-Y | 3 | 3 | 1,2 | 1,4 |
| S-S | 2 | 1 | 2,4 | 0,5 |
| A-G | 2 | 1 | 1,0 | 0,2 |
| Q-R | 2 | 2 | 1,4 | 1,4 |
| P-P | 2 | 1 | 4,7 | 1,5 |

R>K

| | % 1 | % 24 | Enr 1 | Enr 24 |
|---|---|---|---|---|
| Y-Y | 7 | 10 | 2,3 | 3,1 |
| G-Y | 6 | 11 | 1,0 | 1,8 |
| G-G | 5 | 1 | 1,7 | 0,3 |
| K-Y | 5 | 7 | 1,0 | 1,5 |
| S-Y | 5 | 5 | 1,4 | 1,5 |
| N-Y | 3 | 4 | 1,5 | 2,0 |
| K-K | 3 | 0 | 1,6 | 0,2 |
| A-Y | 3 | 4 | 1,2 | 2,0 |
| G-N | 3 | 1 | 1,2 | 0,5 |
| S-S | 3 | 0 | 2,6 | 0,4 |

Y>A

| | % 1 | % 24 | Enr 1 | Enr 24 |
|---|---|---|---|---|
| A-A | 8 | 5 | 1,4 | 0,8 |
| G-G | 5 | 2 | 1,7 | 0,6 |
| R-R | 4 | 2 | 2,2 | 1,2 |
| R-S | 3 | 3 | 1,1 | 1,0 |
| S-S | 3 | 0 | 2,6 | 0,5 |
| Q-R | 2 | 3 | 1,5 | 2,1 |
| P-R | 2 | 2 | 1,2 | 1,3 |
| P-P | 2 | 0 | 5,0 | 1,0 |
| R-T | 2 | 3 | 1,1 | 1,8 |
| P-Q | 2 | 1 | 3,0 | 1,0 |

Enrichment of contacts over sequence background: 5,0 – 0,2

**C**

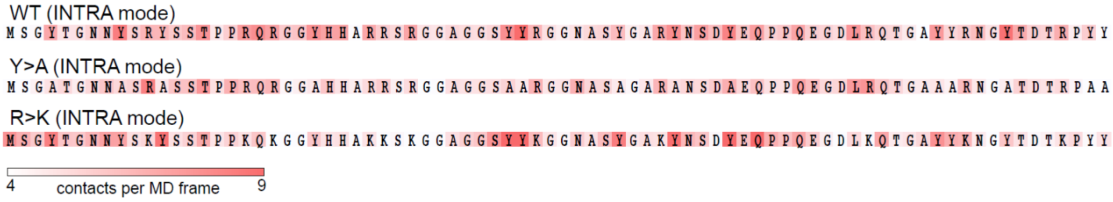


**D**

| contacts | WT | Y>A | R>K |
|---|---|---|---|
| pairwise | 0,96 | 0,96 | 0,98 |
| 2D | -0,02 | -0,02 | -0,03 |

**E**

| modes | WT | R>K | Y>A |
|---|---|---|---|
| INTER repesentative vs average | 0,77 | 0,83 | 0,61 |
| INTER repesentative vs INTRA | 0,05 | 0,18 | 0,20 |
| INTER repesentative vs INTRA-24 | -0,05 | 0,02 | 0,27 |
| INTER average vs INTRA-24 | 0,18 | 0,25 | 0,16 |
| INTRA-24 vs INTRA | 0,71 | 0,75 | 0,88 |

**F**

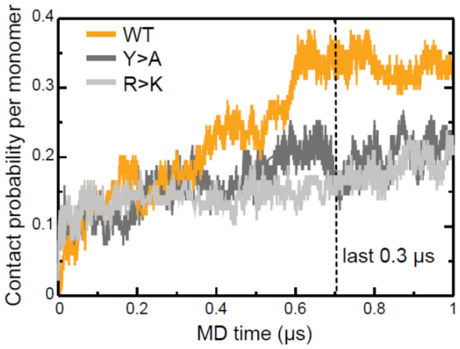


**Figure S3**. **Inter- and intramolecular interactions of Lge1$_{1\text{-}80}$ variants**. (**A**) Top 10 most frequent and enriched pairwise contacts between the residues in different protein molecules in multi-chain simulations (percentages, % 24; enrichment, Enr 24) as compared to single-chain simulations (% 1, Enr 1) for the WT (*left panels*), the R>K (*middle panels*), and the Y>A (*right panels*) Lge1$_{1\text{-}80}$ variants. The enrichments were calculated in comparison to the randomized sequence background according to the scale given on the right. Contact statistics were collected over the last 0.3 µs of MD simulations for two independent single-chain MD runs and all 24 proteins in multi-chain MD runs. (**B**) Top 10 most frequent and enriched pairwise contacts in the single-molecule context as compared to the crowded environment. Details as in A. (**C**) Distributions of statistically defined interaction regions ("stickers") along the protein sequence in the single-chain systems. Protein sequences are colored according to the average contact statistics over the last 0.3 μs in two independent MD runs. (**D**) Pearson correlation coefficients between pairwise contact statistics and 2D pairwise contacts maps obtained over the last 0.3 μs of single- and multi-chain simulations. (**E**) Pearson correlation coefficients calculated from the comparison of statistically defined interaction modes obtained over the last 0.3 μs of single- and multi-chain simulations as shown in panel C and Fig. 3A. (**F**) Time evolution of contact probability per monomer for the WT (*orange line*), Y>A (*dark gray line*), and R>K (*light gray line*) in multi-chain systems.

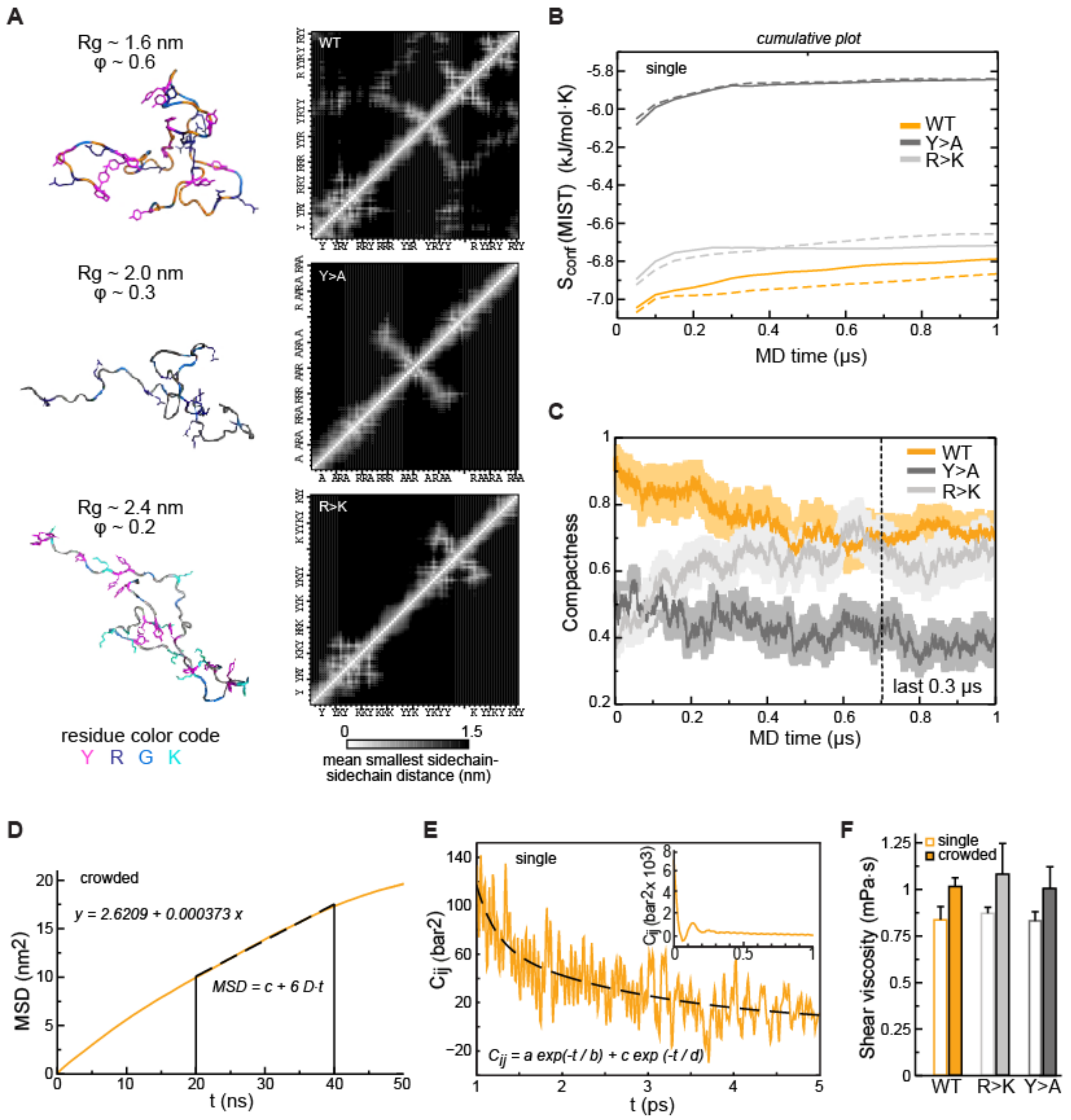


**Figure S4. Conformational behavior and condensate rheology of Lge1$_{1\text{-}80}$ variants**. (**A**) MD conformations of WT (*top panels*), Y>A (*middle panels*), and R>K (*bottom panels*) variants illustrating different levels of compactness with the corresponding inter-residue distance matrices averaged over the 100 ns of a single-chain MD trajectory (contact maps; scale: white– 0 nm; black – 1.5 nm). Proteins are shown in cartoon and sticks representation, and colored in orange (WT), dark gray (Y>A), and gray (R>K) with tyrosine (Y), arginine (R), lysine (K), and glycine (G) residues colored according to the legend given below. (**B**) Convergence of protein configurational entropy ($S_{conf}$) in the single-molecule context. Data for two independent MD replicas are shown with solid and dashed lines, respectively. Cumulative plots were generated using MIST approximation (see Methods) and a 50 ns time step. (**C**) Time evolution of protein compactness for the WT (*orange line*), Y>A (*dark gray line*), and R>K (*light gray line*) in multi-chain systems. The average values over all 24 copies are shown together with the standard errors of the mean. (**D**) Exemplary fitting of an MSD curve in the linear range (20-40 ns) for a protein molecule in multi-chain simulations (shown for Lge1$_{1\text{-}80}$ WT). (**E**) Example of double-exponential fitting of the autocorrelation function for a pressure tensor element in the range of 1-5 ps. Inset: An initial part of the autocorrelation function for a pressure tensor element. Data from the analysis of NVT simulations of a single protein copy (shown for Lge1$_{1\text{-}80}$ WT).

(**F**) Shear viscosity values obtained from the analysis of the pressure tensor autocorrelation functions (see Methods for details) in single-chain context (open bars) and multi-chain context (filled bars). The average values over different pressure tensor elements are shown together. Error bars depict standard deviations.

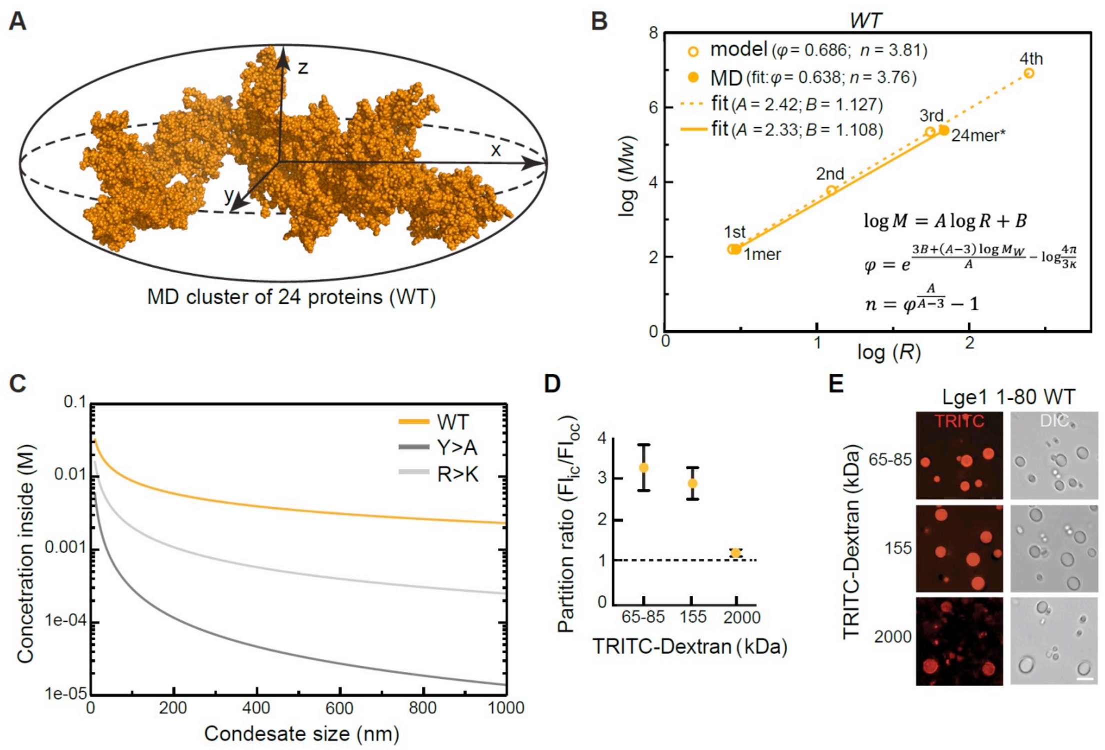


$$\log M = A \log R + B$$

$$\varphi = e^{\frac{3B+(A-3)\log M_W}{A} - \log\frac{4\pi}{3\kappa}}$$

$$n = \varphi^{\frac{A}{A-3}} - 1$$

**Figure S5. Protein clusters and condensate topology of Lge1$_{1\text{-}80}$. (A)** A 24-protein cluster for Lge1$_{1\text{-}80}$ WT fitted to an ellipsoid with radii corresponding to *x*, *y*, *z* components of *Rg*. Protein atoms are shown as spheres. (**B**) Power-law relationship between mass and size of WT protein clusters over four iterations of the fractal model (open circles) with the applied MD-derived valency (*n*) and compactness ($\varphi$) together with the parameters describing the MD clusters of 1 or 24 proteins (filled circles); the corresponding linear regression trends are shown with dashed and solid lines, respectively, with slope (*A*) and intercept (*B*) values indicated in the legend. The $\varphi$ and $n$ values used in the model and given in the legend correspond to the averages over 24 protein copies, while those corresponding to the MD clusters were directly extracted from the slopes and the intercepts of the linear fits performed according to the equations given in the plot (eq. 9,12, 13 in Appendix). All averaging was done over the last 0.3 µs of the multi-chain MD trajectories. The size of the 24-protein MD cluster was estimated as the radius of the sphere with an equivalent apparent volume as the gyration ellipsoid. (**C**) Dependence of protein concentration inside a condensate on its apparent size as predicted by the fractal model. (**D**, **E**) Partitioning of dextran of different sizes into condensates formed by Lge1$_{1\text{-}80}$ WT. Condensates were incubated with TRITC-labelled dextran (final dextran concentration 0.05 mg/ml) for 15 min at 20 °C and imaged by DIC and fluorescence microscopy. Scale bar, 10 µm.

| Name | *Mw*, kDa | *Rg*, nm | *φ* | *n* | *A* | *B* |
|---|---|---|---|---|---|---|
| WT | 9,067 | 1,60 | 0,686 | 3,81 | 2,420 | 1,192 |
| Y>A | 7,778 | 1,97 | 0,368 | 2,31 | 1,635 | 1,249 |
| R>K | 8,759 | 1,68 | 0,612 | 2,08 | 2,088 | 1,293 |

**Figure S6. Parameters used in the fractal model for different Lge1$_{1\text{-}80}$ variants** (related to Figure 5 and 6). Radius of gyration (*Rg*)**,** valency (*n*) and compactness ($\varphi$) averaged over the last 0.3 µs of MD trajectories for all 24 copies. Slope (*A*) and intercept (*B*) of the linear regression for the log *R* vs. log *Mw* plot are derived from the model (eq. 10, 11 in Appendix).
